# Male and female mice achieve equivalent protection against heterologous SARS-CoV-2 infection through divergent immune mechanisms

**DOI:** 10.64898/2026.09.28.755090

**Authors:** Devin Kenney, Mao Matsuo, Andrea Pantelias, Joseph Léger, Ava Kaplan, Faith L. Cerbo, Da-Yuan Chen, Scott R. Adams, Giulia Unali, Douglas C. Wallace, Mohsan Saeed, Anna Belkina, Christelle Harly, Florian Douam

**Affiliations:** Department of Virology, Immunology, and Microbiology, Boston University Chobanian and Avedisian School of Medicine, Boston, MA, USA; Department of Biochemistry and Cell Biology, Boston University Chobanian and Avedisian School of Medicine, Boston MA, USA; National Emerging Infectious Diseases Laboratories, Boston University, Boston, MA, USA; Université de Nantes, INSERM, CNRS, CRCI2NA, Nantes, France; LabEx IGO ‘Immunotherapy, Graft, Oncology’, Nantes, France; Center for Mitochondrial and Epigenomic Medicine, Children’s Hospital of Philadelphia, Philadelphia, PA, USA; Division of Genetic and Genomic Medicine, Department of Pediatrics, Perelman School of Medicine, University of Pennsylvania, Philadelphia, PA, USA; Department of Pathology & Laboratory Medicine, Boston University Chobanian & Avedisian School of Medicine, Boston, MA, USA; Flow Cytometry Core Facility, Boston University Chobanian & Avedisian School of Medicine, Boston, MA, USA

**Author notes:** Corresponding author: Florian Douam.

## Abstract

Biological sex is a major determinant of antiviral immunity, yet how it shapes lung-resident antiviral immunity remains poorly understood. We report that prior mild SARS-CoV-2 BA.1 infection protects mice from lethal heterologous Delta challenge through distinct sex-biased immune mechanisms. Although heterologous protection is clinically equivalent between sexes, T- and B-cell depletions before BA.1 exposure abrogate protection only in males. Depletion with anti-CD4, but not anti-CD8, abolishes protection in both sexes, but co-depletion with anti-CD4/anti-CD8 partially restores protection in females only. Further, females exhibit enhanced IFNγ-associated myeloid inflammation upon secondary Delta infection. Using lymphoid-deficient mice that display female-biased resistance to Delta infection, IFNγ treatment provides partial protection to naïve males against Delta. Female resistance is lost after previous BA.1 exposure, highlighting the importance of lymphoid-mediated inflammatory regulation. Together, equivalent antiviral protection against heterologous respiratory viral infection is governed by sexually dimorphic mechanisms, with significant implications for designing mucosal vaccines and immunotherapies.

## INTRODUCTION

Biological sex is a profound regulator of the immune system^1–4^. At baseline, females have higher numbers of CD4 T cells, B cells, and monocytes/macrophages, and have a higher propensity for hyper-immune activation. Conversely, males have been shown to have higher levels of CD8 T cells and CD4+ Regulatory T cells^5^. Females generally exhibit more robust immune responses during viral infection and mount stronger humoral immunity upon vaccination compared to males^5,6^. In contrast, males show increased susceptibility to severe and fatal disease^5^, while females display greater incidence of post-acute sequelae of viral infection, autoimmune diseases, and chronic inflammation^5,7–11^.

The COVID-19 pandemic and its etiologic agent SARS-CoV-2 exemplified these sex differences, with males exhibiting higher incidence of severe disease and hospitalization and greater risk of lethal infection^12^. Conversely, females were reported to have a higher incidence of post-acute sequelae of COVID-19 (PASC) and more severe symptoms during reinfection^9,13^. Efforts to elucidate the mechanisms underlying sex differences in SARS-CoV-2 immunity have been limited by the largely descriptive nature of clinical studies^14,15^. While *in vivo* mechanistic studies highlighting sex as a core biological factor are mainly in the context of primary infection in immune-naïve animals, immune-experienced mice better reflect human immunobiology^16–18^ and could reveal sex biases in antiviral immunity. SARS-CoV-2 represents a unique tool to conduct such studies due to the broad spectrum of disease severity caused by genetically divergent variants, ranging from the more severe disease associated with the Delta variant to the relatively mild disease caused by the Omicron BA.1 in both humans and mice^19–22^.

Here, we show that BA.1 exposure confers an equivalent, sex-independent protection against Delta-induced lethality in mice that is antigen-specific but not primarily mediated by cross-neutralizing antibodies. However, while both male and female mice require CD4+ cells to promote such a protection, only males require B cells, and protection is partially rescued by CD4/CD8 co-depletion only in females. Further, female protective signatures are associated with more robust type-II interferon and inflammatory responses in the lungs, which are in part linked to the myeloid compartment. Together, our findings illuminate how biological sex imparts distinct mechanisms of pulmonary antiviral immunity to heterologous viral infection yet translate into clinically equivalent protective phenotypes. We also highlight the value of immune-experienced models for uncovering previously unrecognized mechanisms of sex-specific antiviral immunity. Our work has important implications for the design of mucosal vaccines and immunotherapies tailored to maximize efficacy and safety in each sex.

## RESULTS

### Mild Omicron BA.1 infection elicits specific protection against heterologous SARS-CoV-2 infection

Infection of K18-hACE2 transgenic mice with Omicron BA.1 results in attenuated disease compared to previous variants^22,23^, permitting immunological studies after recovery from mild SARS-CoV-2 disease in adult mice. Here, we inoculated 15–26-week-old male and female heterozygous K18-hACE2 mice intranasally with 10^4^ plaque-forming units (pfu) of Omicron BA.1 prior to subsequent infection 30 days later with a dose-equivalent homotypic (BA.1) or heterologous (Delta) variant (**Figure 1A**). Delta was selected as a heterologous challenge strain based on the poor cross-neutralizing activity of BA.1-elicited antibodies against Delta^24,25^ and the high divergence between Delta and BA.1 spike proteins (**Figure S1A**).

**Figure 1.**
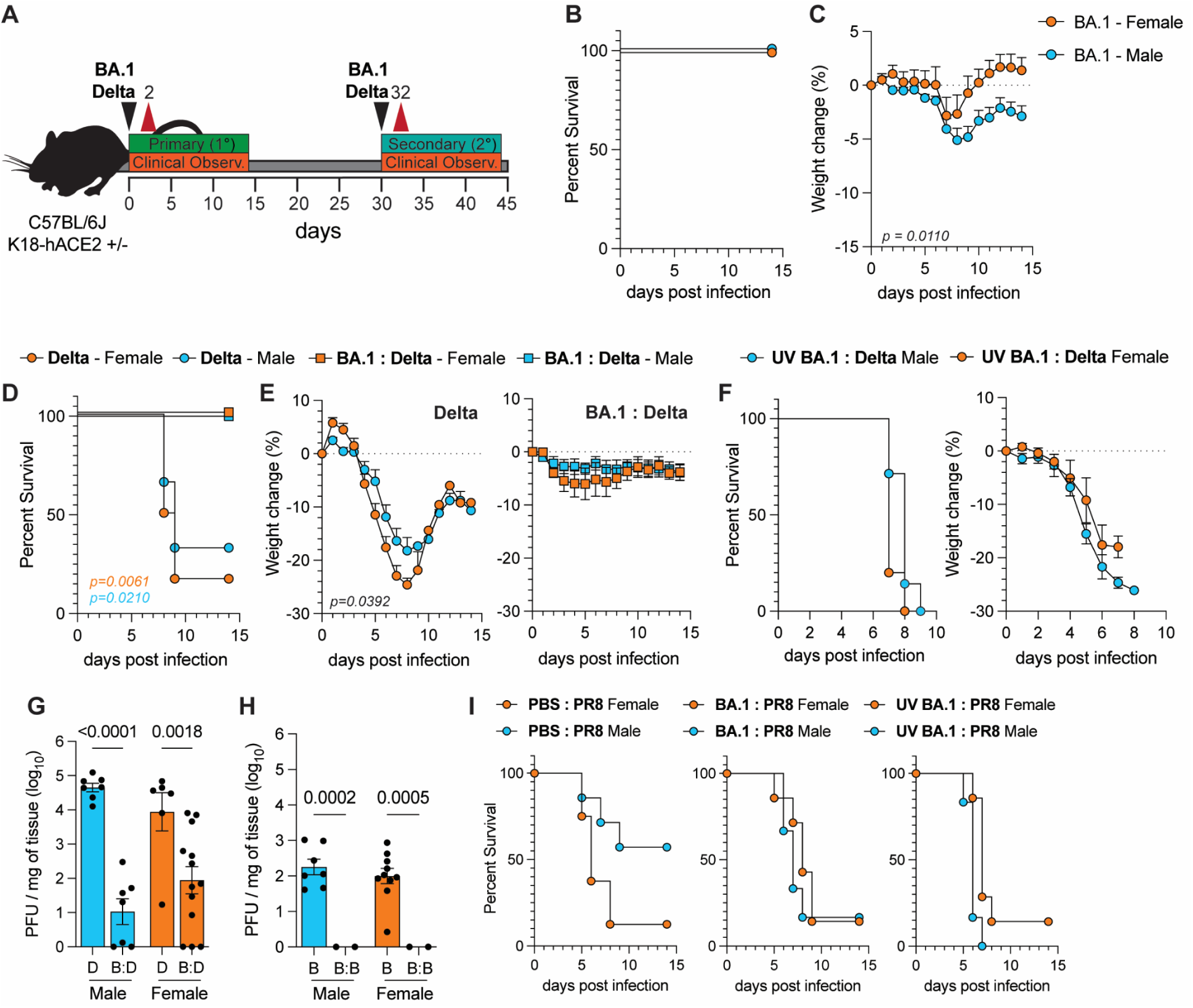
Mild Omicron BA.1 infection elicits specific protection against heterologous SARS-CoV-2 infection. (**A**) Infection scheme for heterologous infection. **(B**) Survival and **(C**) weight change in male (blue, n=17) and female (orange, n = 15) K18-hACE2 mice during primary BA.1 challenge. Mantel-Cox log-rank test. (**D**) Survival and (**E**) weight loss in naive or BA.1**-** exposed male (blue, n=6, n=6) and female (orange, n = 6, n=6) mice after Delta challenge. Mantel-Cox log rank test. **(F)** Survival and weight loss of UV BA.1-exposed male (n=5) and female (n=7) mice after Delta secondary challenge. Mantel-Cox log-rank test. **(G-H)** Viral titers of lung tissue collected at 2 days post-infection from **(G**) Delta primary and secondary infection and (**H**) BA.1 primary and secondary infection. Ordinary two-way ANOVA. Error bars indicate S.E.M. **(I)** Survival of mock (left; n: m= 7, f=8), BA.1-exposed (middle; n: m= 6, f=7), or UV-BA.1 exposed (right; n: m= 6, f=7) male and female mice after secondary infection with 1×10^3^ pfu of Influenza PR8 (H1N1). Mantel-Cox log-rank test.

Lethality was not observed for either sex upon primary BA.1 infection; however, both sexes developed mild disease with peak weight loss occurring between 6 and 8 days post-infection (dpi) (**Figure 1B, C**). Weight loss dynamics were significantly different between males and females, with greater peak weight loss and longer recovery in males **(Figure 1C, S1B**). While Delta primary infection resulted in severe disease and ∼80% lethality, mice previously exposed to BA.1 were protected from Delta lethality, developing only mild disease (**Figure 1D, E**). Notably, females showed more severe weight loss (∼25%) compared to males (∼18%) during primary Delta challenge (**Figure 1E, left panel**), while disease was similar upon secondary challenge (**Figure 1E, right panel**). Homologous secondary infection resulted in minimal disease (**Figure S1C**), and BA.1-exposed mice were fully protected from infection with a chimeric WA-1 virus harboring BA.1 spike (Omi-S)^22^ (**Figure S1D, E**). Ultraviolet (UV) inactivated BA.1 did not protect from Delta secondary challenge, indicating that active replication was required for protection (**Figure 1F**).

No sex bias was observed at 2dpi in the lung of primary BA.1 and Delta-challenged mice, showing that sex does not impact initial susceptibility to infection (**Figure 1G**). Lung titers of BA.1-exposed mice two days after Delta secondary challenge were significantly reduced compared to those from primary Delta-challenged mice. While not significant, BA.1-exposed males showed a greater reduction in Delta viral titer compared to females (**Figure 1G, S1F**). Virus was undetectable in the lungs of BA.1-exposed mice at two days post-secondary homologous BA.1 challenge (**Figure 1H**). Additionally, BA.1-exposed mice were not protected from infection with an antigenically divergent virus, influenza PR8; Notably, disease and lethality were worsened in male mice (**Figure 1I, S1G**), which are normally more resistant than females to PR8 infection^26,27^. Collectively, we show that mild BA.1 infection specifically protects from lethal, heterologous secondary SARS-CoV-2 challenge.

### B cells are necessary for eliciting protective immunity in male but not female mice

To identify the immunological determinants of BA.1-elicited protection, we first assessed the role of neutralizing antibodies (NAb) in serum collected at 30 days post-primary BA.1 infection. Consistent with previous findings^24^, we detected relatively low levels of cross-reactive NAbs targeting Delta **(Figure 2A**). No sex bias or correlation between BA.1 and Delta Nab levels within individual mice was observed (**Figure 2A, S2A**). To determine the *in vivo* protective efficacy of cross-reactive NAbs, 200 µL of serum from BA.1- or PBS-exposed mice was injected intraperitoneally (IP) into naïve K18-hACE2 mice six hours prior to challenge with Delta or Omi-S (**Figure 2B**). Serum from BA.1-exposed mice was insufficient to protect from lethal Delta infection and disease (**Figure 2C**), even when administered intravenously (IV) (**Figure S2B**). However, serum from BA.1-exposed mice conferred complete protection against Omi-S infection (**Figure 2D),** similar to prior BA.1 infection (**Figure S1E**), suggesting homotypic infection is mainly mediated by NAb responses.

**Figure 2.**
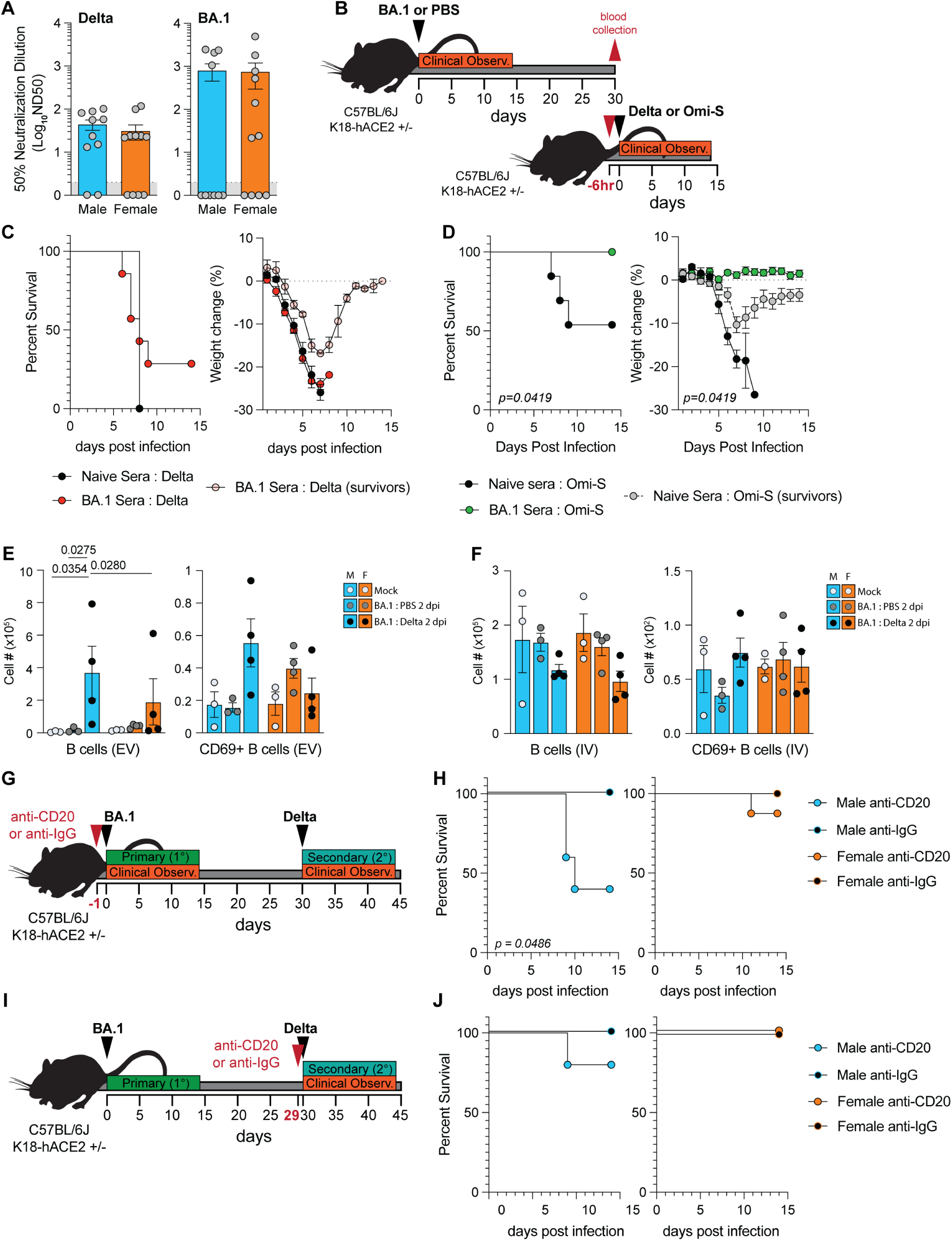
B cells, but not neutralizing antibodies, are primary determinants of protective immunity in male but not female mice. (**A**) Neutralization assay on serum collected at 28-30 days post BA.1 challenge. (**B**) Schematic for serum adoptive transfer experiments. (**C**) Survival (left) and weight change (right) after Delta infection of mice that received intraperitoneal administration of 200 uL of serum from BA.1 (n = 7) or mock mice (n = 7). Survival: Mantel-Cox log-rank test. Weight loss: Multiple t-tests with Bonferroni correction for multiple comparisons. Error bars indicate S.E.M. (**D**) Survival (left) and weight change (right) after Omi-S infection of mice that received intraperitoneal administration of 200 uL of serum from BA.1 (n = 7) or mock mice (n = 13) 6-hours prior to infection. Survival: Mantel-Cox log-rank test. Weight loss: Multiple t-tests with Bonferroni correction for multiple comparisons. Error bars indicate S.E.M. (**E-F**) Flow cytometry of intravascular (IV+) and extravascular (EV) B cell compartments in the lung of mock-infected mice or at 32 days post BA.1 challenge or 2 days post Delta challenge of BA.1-exposed mice. (**E**) Number of EV and (**F**) IV B cells. Two-way ANOVA with Tukey’s correction. Error bars indicate S.E.M. (**G**) Schematic of B-cell depletion prior to BA.1 challenge. (**H**) Survival of anti-CD20 (n: m=5, f=8) or anti-IgG (n: m=5, f=7) depleted mice after secondary Delta infection. Mantel-Cox log-rank test. (**I**) Schematic of B-cell depletion post-BA.1 challenge and prior to Delta secondary challenge. (**J**) Survival of anti-CD20 (n: m=5, f=5) or anti-IgG (n: m=5, f=5) depleted mice after secondary Delta infection. Mantel-Cox log-rank test.

Memory B cells are critical for humoral immunity against heterologous variants of SARS-CoV-2^24,28,29^. Flow cytometry analysis of the lung extravascular (EV) compartment showed significantly more B cells in males 2 days post-secondary challenge compared to females, and a greater number of activated B cells in males, though not significant (**Figure 2E & S3A-D**). However, there was no significant expansion of B cell populations in the lung intravascular (IV) compartment (**Figure 2F**). Consistently, systemic B cell depletion with anti-CD20 (MB20-11) prior to primary BA.1 infection (**Figure 2G, S2C**) significantly reduced protection from secondary Delta infection in males (100% to 40%) but only minimally in females (100% to 87.5%) (**Figure 2H**). Interestingly, mice that survived infection did not experience severe disease, implying that those that lost protection succumbed to disease (**Figure S2D**). Systemic B cell depletion prior to Delta secondary challenge (**Figure 2I**) did not significantly increase lethality or disease in male or female mice (**Figure 2J, S2E**). Importantly, MB20-11 antibody depletes B cells in circulation via liver and splenic macrophages^30,31^, making a post-BA.1 challenge depletion unlikely to deplete lung resident memory B cells. Collectively, these findings suggest that loss of B cells during BA.1 exposure abrogates heterologous protection against Delta infection in males, but not females.

### T cells play divergent roles in male and female immunity during secondary infection

In addition to B cells, analysis of the EV lung compartment showed a larger increase in memory CD4 and CD8 T cells after secondary Delta infection in males compared to females (**Figure 3A, B & S3A, B**), whereas lung IV T cell populations were not increased in either sex (**Figure 3C, D & S3C, D**). T cell depletion prior to secondary Beta (B.1.351) infection in BA.1-exposed females has been shown not to impede protection^32^. However, T cell functionality differs between sexes^33^, and the contribution of T cells to protection against secondary infection remains to be investigated in males. Therefore, we systemically depleted CD3+ cells prior to and during primary BA.1 challenge using a CD3e F(ab’)_­_antibody (clone 30-F11 Fab) (**Figure 3E**), which, unlike anti-CD3e treatment, depletes only antigen-activated T cells and does not induce T cell activation^34^ and associated disease (**Figure S4A,B**). Anti-CD3e F(ab’)_­_treatment resulted in 100% lethality in BA.1-exposed males upon Delta challenge, compared to ∼29% lethality in anti-IgG F(ab’)_­_control-treated mice (**Figure 3F)**. Comparatively, no significant increase in lethality was seen in CD3e F(ab’)_­_-depleted females (**Figure 3F**), and weight loss was not observed in mice that survived infection (**Figure S4C**). Importantly, sex bias was not due to differential depletion efficiencies (**Figure S4D**). T-cell depletion following BA.1 primary challenge (**Figure S4E**) did not affect the survival or weight loss of either sex (**Figure 3G, S4F**) and adoptive transfer of splenic T cells from BA.1-exposed mice into naïve mice did not confer protection in either sex post-secondary Delta challenge (**Figure S4G-I**), indicating that circulating memory T cells are neither necessary nor sufficient for protection. These findings support a critical role of T cells in male, but not female, protection.

**Figure 3.**
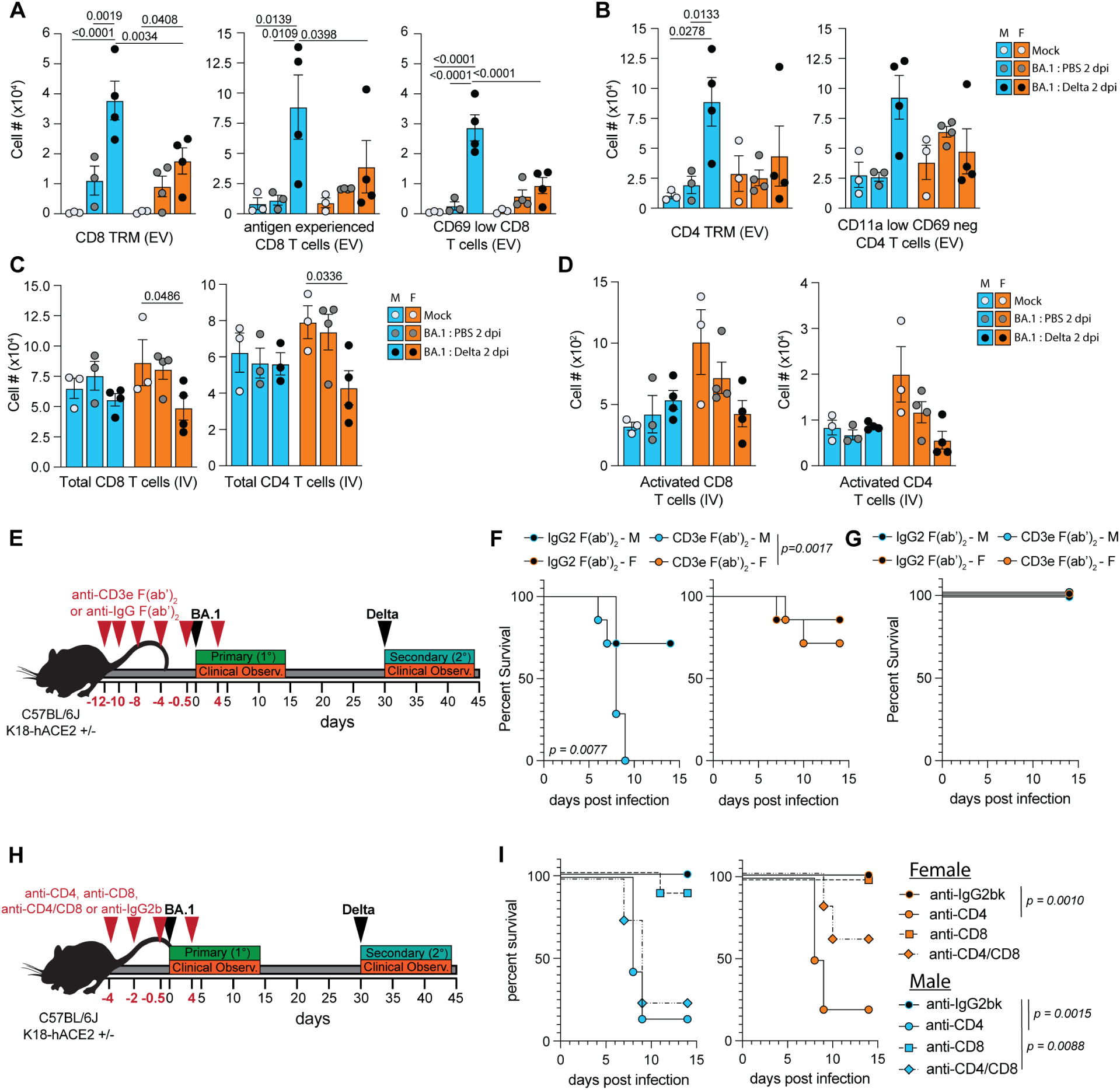
T cells play divergent roles in male and female immunity during secondary infection. (**A-D**) Flow cytometry of intravascular (IV) and extravascular (EV) T cell compartments in the lung of mock-infected mice or at 32 days post-BA.1 challenge or 2 days post-Delta challenge of BA.1-exposed mice. (**A**) Number of EV CD8 T cell populations, (**B**) EV CD4 T cell populations, (**C**) IV T cell populations, and (**D**) activated IV T cell populations. Ordinary two-way ANOVA with Tukey’s correction. Error bars indicate S.E.M. (**E**) Schematic of T cell depletion with anti-CD3e F(ab’)_­­_prior to and during BA.1 primary infection. (**F**) Survival curve for Delta secondary infection of mice depleted with anti-CD3e F(ab’)_­_(n: m=7, f=7) or anti-IgG F(ab’)_­_(n: m=7, f=7) prior to and during BA.1 primary infection. Mantel-Cox log-rank test. (**G**) Survival after Delta challenge of mice depleted with anti-CD3e F(ab’)_­_(n: m=3, f=3) or anti-IgG F(ab’)_­_(n: m=3, f=3) after BA.1 primary infection and during Delta secondary infection. Mantel-Cox log-rank test. (**H**) Schematic for depletion with anti-CD4, anti-CD8, or anti-CD4/anti-CD8 or anti-IgG2bk prior to and during primary BA.1 challenge. (**I**) Survival after Delta challenge of mice depleted with anti-CD4 (n: f=10, f=7), anti-CD8 (n: m=8, f=9), or anti-CD4/anti-CD8 (n: m=4, f=5) or anti-IgG2bk (n: m=7, f=8) prior to and during BA.1 primary infection. Mantel-Cox log-rank test.

We next depleted CD4- and CD8-expressing cells prior to BA.1 infection (**Figure 3H, S4J**). CD8 depletion had no impact on protection in female BA.1-exposed mice and minimal impact in males; however, CD4 depletion resulted in significant lethality and disease in both male (84%) and female (80%) mice (**Figure 3I, S4K**). Given that females remained protected after CD3 F(ab’)_­_depletion, we asked whether CD8 cells might mediate pathology in females in the absence of CD4 cells. Co-depletion with anti-CD4/CD8 (**Figure 3H)** increased survival to 60% compared to 20% in CD4-depleted female mice (**Figure 3I, S4K**). However, consistent with CD3 F(ab’)_­_depletion, co-depletion in males did not reduce lethality (**Figure 3I**, **S4K**). Together, our findings suggest that CD4 T cells are critical for both male and female heterologous protection, but most likely exert their protective role in distinct, sexually dimorphic ways.

### Females exhibit robust type II interferon and inflammatory responses upon reinfection

Female mice exhibit stronger innate responses, partly due to higher expression of X-linked immune genes that escape Xist-mediated silencing (such as *TLR7, CXCR3* and *CD40L*^35^), as well as the immunostimulatory effects of 17β-estradiol and the immunosuppressive effects of androgens^5,36,37^. Serum cytokine analysis at 2 days post-Delta secondary challenge revealed a significant upregulation of CXCL10 (IP10) and IFNγ in females and IFNγ in males (**Figure 4A**). While serum IFNγ upregulation was not sexually dimorphic, RT-qPCR revealed stronger *Ifng* upregulation in the lung of female mice (**Figure 4B**). Significant upregulation of serum IFNγ was not observed at 2 dpi during primary Delta infection or in BA.1-exposed mice prior to rechallenge (**Figure S4L, M**).

**Figure 4.**
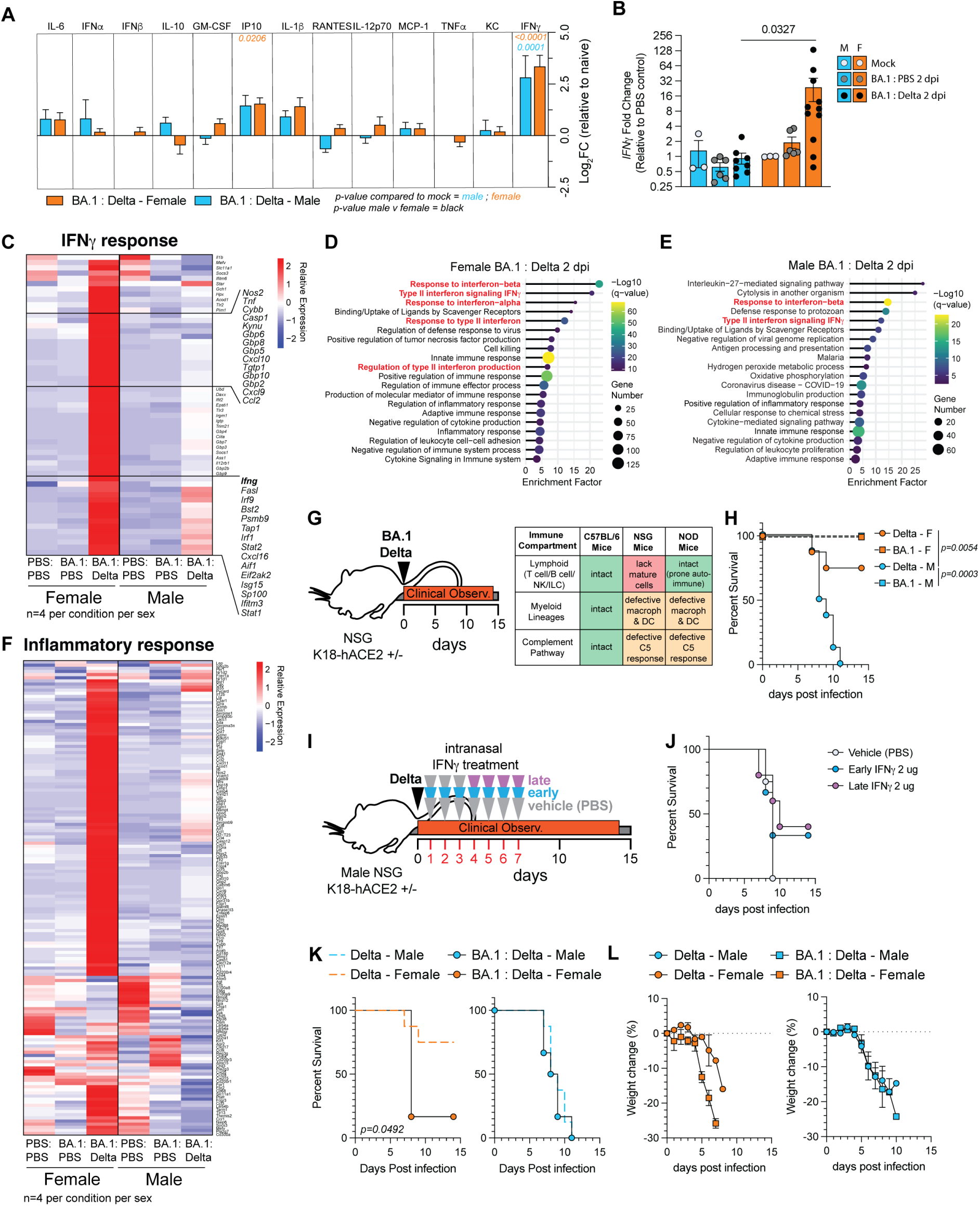
Female mice exhibit stronger inflammatory and type-II IFN responses early after Delta reinfection. (**A**) BioLegend LegendPlex 13-plex antiviral cytokine quantification from serum collected two days post-Delta secondary challenge. Log_­_Fold-change normalized to serum from mock. m=3, f=9. Multiple t-tests with Bonferroni correction for multiple comparisons. Error bars indicate S.E.M. (**B**) RT-qPCR analysis of *Ifng* on lung tissues collected at 32 days post primary BA.1 infection (n: m=6, f=6) and 2 days post delta secondary infection (n: m=8, f=11) or mock mice (n: m=3, f=3). Log_­_Fold-change normalized to mock. Multiple t-tests with Bonferroni correction for multiple comparisons. Error bars indicate S.E.M. **(C-F**) Bulk RNA-sequencing analysis from lung tissues collected at 32 days post primary BA.1 infection and 2 days post delta secondary infection or mock mice (n: m=4, f=4). (**C**) Heatmap indicating relative expression of genes associated with IFNγ response. **(D-E)** GO term analysis performed on significantly upregulated genes in lungs of female (**D**) and male (**E**) at 2 days post-secondary delta challenge. (**F**) Heatmap indicating relative expression of genes associated with inflammatory responses. (**G**) Schematic of NSG K18-hACE2 mouse infection. (**H**) Survival of NSG-K18-hACE2 mice infected with 10^4^ pfu of SARS-CoV-2 Delta. Mantel-Cox log-rank test. (**I**) Schematic of IFNγ treatment in NSG-hACE2 mice. NSG-hACE2 mice were intranasally administered 2 µg daily starting at 1 dpi (early) or 4 dpi (late), or with PBS, prior to primary Delta challenge. (**J**) Survival of NSG-K18-hACE2 mice treated with IFNγ and infected with 10^4^ pfu of SARS-CoV-2 Delta. Mantel-Cox log-rank test. (**K**) Survival of BA.1-exposed NSG-K18-hACE2 mice infected with 10^4^ pfu of SARS-CoV-2 Delta. Mantel-Cox log-rank test. (**L**) Weight loss in naïve or BA.1-exposed mice that died after infection with 10^4^ pfu of SARS-CoV-2 Delta.

Bulk RNA-sequencing of lung homogenates supported a stronger IFNγ-associated response in BA.1-exposed females compared to males upon secondary Delta challenge (**Figure 4C**), which included upregulation of myeloid and macrophage-associated inflammatory genes (e.g, Gbp proteins, and *Ciita*) and other pro-inflammatory mediators (e.g., *Cxcl10, Cxcl9*). IFNγ-associated and inflammatory pathways were also broadly enriched in females compared to males (**Figure 4D-F**). Notably, lymphopenia during SARS-CoV-2 infection has been correlated with excessive inflammation^38^, and BA.1-exposed female mice displayed a mild lymphopenia in the IV immune compartment after Delta challenge, which was more prominent compared to males (**Figure 2F, 3C)**. In males, the decrease in IV B cells and CD4 T cells correlated with a significant increase in EV cells, though this was not the case for females (**Figure 2E, F & 3B-D**).

Collectively, protection of BA.1-exposed females against secondary Delta challenge is associated with a robust type-II IFN-associated, signature of myeloid activation, and overall inflammatory responses.

### BA.1 priming exacerbates Delta-induced disease in female, but not male, lymphoid-deficient NSG-hACE2 mice

Given these findings, we hypothesized that female protection during Delta secondary infection is 1) driven in part by IFNγ-regulated myeloid responses and 2) reliant on CD4 T cells to help control myeloid inflammation. To test this hypothesis, we utilized NSG-K18-hACE2 mice that lack mature lymphocytes but maintain a myeloid compartment, albeit with some functional deficits^39–41^ (**Figure 4G**). Notably, female non-obese diabetic (NOD) mice expressing the K18-hACE2 cassette, which represent the background strain of NSG-K18-hACE2 mice, are more resistant to lethal SARS-CoV-2 infection compared to males^42^. Consistently, Delta infection of NSG-K18-hACE2 mice resulted in ∼25% lethality in females and 100% lethality in males, while BA.1 infection remained non-lethal regardless of sex (**Figure 4H**). To determine if IFNγ stimulation of a myeloid-only immune compartment is sufficient to rescue male lethality, we administered IFNγ intranasally starting at 1 day (early) or 4 days (late) post-Delta infection (**Figure 4I**). Both early and late treatments partially rescued male lethality after Delta infection (**Figure 4J**). Notably, while BA.1 primary infection did not prevent Delta-induced lethality in males, BA.1-exposed females displayed increased lethality (83%) following Delta secondary challenge (**Figure 4K**). Additionally, BA.1-exposed females that succumbed to secondary Delta infection showed greater weight loss compared to lethal primary Delta infection, unlike males (**Figure 4L**). Together, we show that IFNγ stimulation of a myeloid-only immune compartment can partially rescue lethal Delta infection in males, while complete priming of the myeloid compartment with BA.1 in the absence of a lymphoid compartment abrogates the intrinsic protection seen in female NSG-K18-hACE2 mice.

### Female-bias loss of heterologous protection in aged mice

Sex-biased immune functions become increasingly divergent as individuals age^4,43^, most notably in CD8 T cells and inflammatory monocytes^4,44^. Therefore, we hypothesized that these sex-biased immune phenotypes may be impacted in aged mice (85-95 weeks). Notably, while aged C57BL/6J K18-hACE2 males were more susceptible to weight loss during primary BA.1 infection than aged females **(Figure S4N, O**), BA.1-exposed aged females were more susceptible to loss of heterologous protection against Delta than BA.1-exposed aged males (**Figure S4P, Q**). By demonstrating that equivalent clinical protection against heterologous infection across sexes is age-dependent, these findings provide orthogonal validation of the critical role of biological sex in heterologous antiviral immunity in mice.

## DISCUSSION

Collectively, our study shows that after an initial respiratory viral infection, male and female mice leverage distinct immune mechanisms to protect from a secondary, heterologous infection that evades antibody responses. We found that CD4 T cells and B cells, but not CD8 T cells, were required to promote a protective lung environment in males against subsequent heterologous Delta infection following an initial mild BA.1 exposure. This was associated with higher numbers of extravascular activated B cells and memory CD4+ and CD8+ T cells in the lungs of males than females. While female heterologous protection was only slightly abrogated upon CD3 or B cell depletion, protection was lost upon CD4 depletion. Co-depletion of CD4 and CD8 cells partially rescued CD4 lethality in females but not males. Female BA.1-exposed mice showed higher IFNγ and myeloid-associated inflammatory responses upon secondary Delta infection, suggesting female-biased hyperinflammatory responses. Notably, aged mice exhibited a more extensive loss of heterologous protection in females than males, further validating that sexually dimorphic immune mechanisms govern heterologous protection.

Our findings strongly align with evidence that anti-CD3 F(ab’)_­_does not impact T regulatory cells^45^, thus allowing for some regulation of inflammation, while this population is missing in NSG-hACE2 mice and upon CD4 depletion. Loss of protection, either in males or females, occurred only when depletion was performed systemically prior to and/or during BA.1 initial exposure, but not prior to and/or during secondary Delta infection, highlighting that lung-resident immune populations are sufficient to elicit protection. The absence of protection against Influenza PR8 virus in BA.1-exposed mice demonstrates an antigen-specific response and, therefore, likely extends beyond lung-resident trained innate immune mechanisms.

We propose a model in which lung-resident immune remodeling during mild BA.1 infection drives sexually dimorphic responses during a subsequent, heterologous SARS-CoV-2 infection. CD4 T cells are an essential component of the immune response upon primary infection for effective heterologous protection in both males and females, though via distinctive pathways. In males, consistent with higher numbers of extravascular activated B cells and CD4+ T cells in males, CD4 T cells engage in crosstalk with B cells to establish a protective lung environment that supports robust antigen-specific responses upon heterologous challenge. In contrast, CD4 T cells seem to play a critical role in the regulation of IFNγ-associated inflammatory responses, likely driven by T cell-myeloid crosstalk, in BA.1-exposed females.

These data also provide insight into the potential mechanisms that cause higher incidence of PASC in women^9,46^, especially in non-hospitalized patients^7^. An IFNγ-mediated inflammatory feedback loop between CD8 T cells, fibroblasts, and macrophages has been proposed as a major contributor to PASC and other post-acute pulmonary viral diseases^47,48^. Elevated IFNγ-associated inflammation and deleterious CD8 T cell responses observed in females suggest a mechanistic rationale for why women are more prone to PASC and raise the question of what the sex-linked factors driving this phenotype are. This work also opens major avenues to leverage immune-experienced mouse models to mechanistically dissect upstream and downstream regulators of lung-resident sexually dimorphic immunity during respiratory viral infection, including hormones and X-linked genetic factors, and to define how these mechanisms are altered during aging. Importantly, given that the historical underrepresentation of women in clinical trials has contributed to female biases in vaccine- and treatment-associated adverse effects^49,50^, our findings identify cellular immune mechanisms that should be considered when designing vaccines and immunotherapies for pulmonary viral infections to minimize female-biased adverse effects. Finally, our work raises fundamental questions regarding the evolutionary pressures that drive males and females to adopt distinct, yet clinically convergent, immune mechanisms to fight successive respiratory infections, and whether such sex-specific adaptations extend beyond the lung.

### Limitations of the study

We acknowledge limitations to the current study. First, despite BA.1 exposure, the immune compartment of the mouse lung remains relatively naïve compared to humans. Our study is also limited to a single inbred mouse strain, except for the final studies in NSG mice. Validating these findings in other genetic backgrounds, in more complex immune-experienced animals (e.g., pet store mice), and across a wider SARS-CoV-2 antigenic landscape would enrich our findings. Also, CD4 and CD8 can be expressed on other cell types (e.g., cDC1 and NK); therefore, the impact of these cells on our protection phenotype cannot be excluded. Despite these limitations, our study provides new insight into the impact of biological sex on pulmonary immunobiology during viral infection, underscoring the importance of considering biological sex.

## ACKNOWLEDGMENTS

This work was supported in part by Boston University/NEIDL start-up funds (to F.D. and M.S.), a Peter Paul Career Development Professorship from Boston University (to F.D.), Clinical and Translational Science Awards (grant UL1 TR001430) from the National Center for Advancing Translational Sciences of the National Institutes of Health (to F.D.), and the Affinity Research Collaborative on ‘Respiratory Viruses: A Focus on COVID-19’ at the Evans Center for Interdisciplinary Biomedical Research at Boston University Chobanian & Avedisian School (to F.D.). D.K. received support from an NIH T32 Immunology Training Program (T32AI007309) and by a Hartwell Postdoctoral Fellowship awarded by The Hartwell Foundation (55212221). D.C.W. received support from NIH AG078814, CA259635, and DOD W81XMH2110128 PR202887. This work was also supported by the Boston University Flow Cytometry Core Facility. We thank the Boston University Animal Science Center, and the NEIDL animal core staff for their outstanding support. We also thank all the Douam Lab, Saeed Lab, and NEIDL members, and members of the Department of Virology, Immunology and Microbiology and Department of Biochemistry & Cell Biology at Boston University for their constant support and advice. Some figures were created using Biorender.com. We used Grammarly (Grammarly Inc.) to assist with grammar correction and sentence structure optimization during manuscript preparation.

## AUTHOR CONTRIBUTIONS

**Devin Kenney:** Conceptualization, Methodology, Investigation, Formal Analysis, Data Curation, Visualization, Writing – Original Draft, Writing – Review & Editing; **Mao Matsuo**: Investigation, Writing – Review & Editing; **Andrea Pantelias**: Investigation, Writing – Review & Editing; **Joseph Léger**: Formal analysis, Visualization, Writing – Review & Editing; **Ava Kaplan:** Investigation, Writing – Review & Editing; **Faith L. Cerbo**: Investigation, Writing – Review & Editing; **Da-Yuan Chen**: Resources, Writing – Review & Editing; **Scott R. Adams**: Investigation, Writing – Review & Editing; **Giulia Unali**: Investigation, Writing – Review & Editing; **Douglas C. Wallace**: Resources, Writing – Review & Editing; **Mohsan Saeed**: Resources, Writing – Review & Editing; **Anna Belkina**: Formal analysis, Resources, Writing – Review & Editing; **Christelle Harly**: Formal analysis, Writing – Review & Editing; **Florian Douam**: Conceptualization, Methodology, Funding Acquisition, Project Administration, Resources, Supervision, Writing – Original Draft, Writing – Review & Editing.

## DECLARATION OF INTERESTS

The authors declare no conflict of interest.

## MATERIALS AND METHODS

A detailed description of materials and methods is available in Supplementary Materials.

### Institutional approvals

All animal experiments described in this study were performed in accordance with protocols that were reviewed and approved by the Institutional Animal Care and Use and Committee of Boston University (PROTO202000020; Approval date: 06/15/2020). All mice were maintained in facilities accredited by the Association for the Assessment and Accreditation of Laboratory Animal Care (AAALAC). All replication-competent SARS-CoV-2 experiments were performed in a biosafety level 3 laboratory (BSL-3) at the Boston University National Emerging Infectious Diseases Laboratories (NEIDL).

### Mouse infections

Mice were intranasally inoculated with 1×10^4^ pfu SARS-CoV-2 variants (indicated in text for each experiment) suspended in 50 µL of 1x PBS. For all infections, the 50 µL was split between two nostrils. Secondary infections were carried out 30 days post-primary infection, except for UV-inactivated BA.1 experiments and PR8 secondary challenge, which were performed 40 days post-primary infection. For PR8 infections, mice were infected with 1×10^3^ pfu of influenza PR8 (H1N1) in 50 µL of 1x PBS. All infections were performed while mice were under anesthesia with 1-3% isoflurane.

### Statistical Analysis

For all survival curve comparisons, a Log-rank (Mantel-Cox) test was performed, with p-values of <0.05 being statistically significant. For viral quantification (pfu), flow cytometry, cytokine, neutralizing antibody, and RT-qPCR analyses were performed using One-way or Two-way analysis of variance (ANOVA) tests, as described in the figure legends. All error bars are indicated as mean±standard error of the mean (SEM). For all graphs and data, a p-value of <0.05 was considered significant and indicated on figures and all non-significant values are not indicated. All statistical tests and graphical depictions of results were performed using GraphPad Prism version 10.1.1 software (GraphPad Software, La Jolla, CA). All p-values are indicated on graphs.

### Data availability

The raw bulk RNA-sequencing data are available under GEO accession number GSE345444.

## Notes

### Competing Interest Statement

The authors have declared no competing interest.

### Summary of Updates

We are providing a revised version of the manuscript with improved formatting and corrected typographical errors.

## REFERENCES

1. Edwards, K., Merrill, S.M., Konwar, C., Jude, M.S., Zhuang, B.C., Meijer, M., Navarro-Delgado, E., MacIsaac, J.L., Mandhane, P., Simons, E., et al. (2025). Biological sex impacts immune cell proportions and epigenetic profiles in the developing pediatric immune system. Commun Biol 8, 1447. 10.1038/s42003-025-08844-9.

2. Gheitasi, R., Baumgart, S., Roell, D., Rose, N., Watzl, C., Dudziak, D., Andreas, N., Makarewicz, O., Drube, S., Schnizer, C., et al. (2025). Age- and sex-associated differences in immune cell populations. iScience 28, 113092. 10.1016/j.isci.2025.113092.

3. Huang, Z., Chen, B., Liu, X., Li, H., Xie, L., Gao, Y., Duan, R., Li, Z., Zhang, J., Zheng, Y., and Su, W. (2021). Effects of sex and aging on the immune cell landscape as assessed by single-cell transcriptomic analysis. Proc Natl Acad Sci U S A 118. 10.1073/pnas.2023216118.

4. Sopena-Rios, M., Ripoll-Cladellas, A., Omidi, F., Ballouz, S., Alquicira-Hernandez, J., Oelen, R., Hewitt, A.W., Franke, L., van der Wijst, M.G.P., Powell, J.E., and Mele, M. (2026). Single-cell analysis of the human immune system reveals sex-specific dynamics of immunosenescence. Nat Aging 6, 932–949. 10.1038/s43587-026-01099-x.

5. Sharma, S., Gibbons, A., and Saphire, E.O. (2025). Sex differences in tissue-specific immunity and immunology. Science 389, 599–603. 10.1126/science.adx4381.

6. Klein, S.L., and Flanagan, K.L. (2016). Sex differences in immune responses. Nat Rev Immunol 16, 626–638. 10.1038/nri.2016.90.

7. Gebhard, C.E., Sutsch, C., Gebert, P., Gysi, B., Bengs, S., Todorov, A., Deforth, M., Buehler, P.K., Meisel, A., Schuepbach, R.A., et al. (2024). Impact of sex and gender on post-COVID-19 syndrome, Switzerland, 2020. Euro Surveill 29. 10.2807/1560-7917.ES.2024.29.2.2300200.

8. Silva, J., and Iwasaki, A. (2024). Sex differences in postacute infection syndromes. Sci Transl Med 16, eado2102. 10.1126/scitranslmed.ado2102.

9. Shah, D.P., Thaweethai, T., Karlson, E.W., Bonilla, H., Horne, B.D., Mullington, J.M., Wisnivesky, J.P., Hornig, M., Shinnick, D.J., Klein, J.D., et al. (2025). Sex Differences in Long COVID. JAMA Netw Open 8, e2455430. 10.1001/jamanetworkopen.2024.55430.

10. Chamekh, M., Deny, M., Romano, M., Lefevre, N., Corazza, F., Duchateau, J., and Casimir, G. (2017). Differential Susceptibility to Infectious Respiratory Diseases between Males and Females Linked to Sex-Specific Innate Immune Inflammatory Response. Front Immunol 8, 1806. 10.3389/fimmu.2017.01806.

11. So, J., Tai, A.K., Lichtenstein, A.H., Wu, D., and Lamon-Fava, S. (2021). Sexual dimorphism of monocyte transcriptome in individuals with chronic low-grade inflammation. Biol Sex Differ 12, 43. 10.1186/s13293-021-00387-y.

12. Peckham, H., de Gruijter, N.M., Raine, C., Radziszewska, A., Ciurtin, C., Wedderburn, L.R., Rosser, E.C., Webb, K., and Deakin, C.T. (2020). Male sex identified by global COVID-19 meta-analysis as a risk factor for death and ITU admission. Nat Commun 11, 6317. 10.1038/s41467-020-19741-6.

13. Bai, F., Tomasoni, D., Falcinella, C., Barbanotti, D., Castoldi, R., Mule, G., Augello, M., Mondatore, D., Allegrini, M., Cona, A., et al. (2022). Female gender is associated with long COVID syndrome: a prospective cohort study. Clin Microbiol Infect 28, 611 e619–611 e616. 10.1016/j.cmi.2021.11.002.

14. Sauerwald, N., Zhang, Z., Ramos, I., Nair, V.D., Soares-Schanoski, A., Ge, Y., Mao, W., Alshammary, H., Gonzalez-Reiche, A.S., van de Guchte, A., et al. (2022). Pre-infection antiviral innate immunity contributes to sex differences in SARS-CoV-2 infection. Cell Syst 13, 924–931 e924. 10.1016/j.cels.2022.10.005.

15. Takahashi, T., Ellingson, M.K., Wong, P., Israelow, B., Lucas, C., Klein, J., Silva, J., Mao, T., Oh, J.E., Tokuyama, M., et al. (2020). Sex differences in immune responses that underlie COVID-19 disease outcomes. Nature 588, 315–320. 10.1038/s41586-020-2700-3.

16. Beura, L.K., Hamilton, S.E., Bi, K., Schenkel, J.M., Odumade, O.A., Casey, K.A., Thompson, E.A., Fraser, K.A., Rosato, P.C., Filali-Mouhim, A., et al. (2016). Normalizing the environment recapitulates adult human immune traits in laboratory mice. Nature 532, 512–516. 10.1038/nature17655.

17. Reese, A.T., Chadaideh, K.S., Diggins, C.E., Schell, L.D., Beckel, M., Callahan, P., Ryan, R., Emery Thompson, M., and Carmody, R.N. (2021). Effects of domestication on the gut microbiota parallel those of human industrialization. Elife 10. 10.7554/eLife.60197.

18. Erickson, S., Lauring, B., Cullen, J., and Medzhitov, R. (2026). Environmentally driven immune imprinting protects against allergy. Nature 650, 987–996. 10.1038/s41586-025-10001-5.

19. Lin, L., Liu, Y., Tang, X., and He, D. (2021). The Disease Severity and Clinical Outcomes of the SARS-CoV-2 Variants of Concern. Front Public Health 9, 775224. 10.3389/fpubh.2021.775224.

20. Halfmann, P.J., Iida, S., Iwatsuki-Horimoto, K., Maemura, T., Kiso, M., Scheaffer, S.M., Darling, T.L., Joshi, A., Loeber, S., Singh, G., et al. (2022). SARS-CoV-2 Omicron virus causes attenuated disease in mice and hamsters. Nature 603, 687–692. 10.1038/s41586-022-04441-6.

21. van Doremalen, N., Singh, M., Saturday, T.A., Yinda, C.K., Perez-Perez, L., Bohler, W.F., Weishampel, Z.A., Lewis, M., Schulz, J.E., Williamson, B.N., et al. (2022). SARS-CoV-2 Omicron BA.1 and BA.2 are attenuated in rhesus macaques as compared to Delta. Sci Adv 8, eade1860. 10.1126/sciadv.ade1860.

22. Chen, D.Y., Chin, C.V., Kenney, D., Tavares, A.H., Khan, N., Conway, H.L., Liu, G., Choudhary, M.C., Gertje, H.P., O’Connell, A.K., et al. (2023). Spike and nsp6 are key determinants of SARS-CoV-2 Omicron BA.1 attenuation. Nature 615, 143–150. 10.1038/s41586-023-05697-2.

23. Diamond, M., Halfmann, P., Maemura, T., Iwatsuki-Horimoto, K., Iida, S., Kiso, M., Scheaffer, S., Darling, T., Joshi, A., Loeber, S., et al. (2021). The SARS-CoV-2 B.1.1.529 Omicron virus causes attenuated infection and disease in mice and hamsters. Res Sq. 10.21203/rs.3.rs-1211792/v1.

24. Suryawanshi, R.K., Chen, I.P., Ma, T., Syed, A.M., Brazer, N., Saldhi, P., Simoneau, C.R., Ciling, A., Khalid, M.M., Sreekumar, B., et al. (2022). Limited cross-variant immunity from SARS-CoV-2 Omicron without vaccination. Nature 607, 351–355. 10.1038/s41586-022-04865-0.

25. Ma, J., Liu, X., Zhou, M., Chen, P., Chen, R., Wang, J., Zhu, H., Wu, K., Ye, J., Zhang, Y., et al. (2023). A Heterologous Challenge Rescues the Attenuated Immunogenicity of SARS-CoV-2 Omicron BA.1 Variant in Syrian Hamster Model. J Virol 97, e0168422. 10.1128/jvi.01684-22.

26. Sabikunnahar, B., Lahue, K.G., Asarian, L., Fang, Q., McGill, M.M., Haynes, L., Teuscher, C., and Krementsov, D.N. (2022). Sex differences in susceptibility to influenza A virus infection depend on host genotype. PLoS One 17, e0273050. 10.1371/journal.pone.0273050.

27. Klein, S.L., Hodgson, A., and Robinson, D.P. (2012). Mechanisms of sex disparities in influenza pathogenesis. J Leukoc Biol 92, 67–73. 10.1189/jlb.0811427.

28. Kaku, C.I., Bergeron, A.J., Ahlm, C., Normark, J., Sakharkar, M., Forsell, M.N.E., and Walker, L.M. (2022). Recall of preexisting cross-reactive B cell memory after Omicron BA.1 breakthrough infection. Sci Immunol 7, eabq3511. 10.1126/sciimmunol.abq3511.

29. Chu, Q., Li, K., He, Q., Ren, L., Wang, J., Wang, S., Liu, X., Liu, Y., He, J., Li, D., and Shao, Y. (2025). Efficient boosting of Omicron-reactive memory B cells after breakthrough infection protects from repeated exposure. iScience 28, 112278. 10.1016/j.isci.2025.112278.

30. Uchida, J., Hamaguchi, Y., Oliver, J.A., Ravetch, J.V., Poe, J.C., Haas, K.M., and Tedder, T.F. (2004). The innate mononuclear phagocyte network depletes B lymphocytes through Fc receptor-dependent mechanisms during anti-CD20 antibody immunotherapy. J Exp Med 199, 1659–1669. 10.1084/jem.20040119.

31. Montalvao, F., Garcia, Z., Celli, S., Breart, B., Deguine, J., Van Rooijen, N., and Bousso, P. (2013). The mechanism of anti-CD20-mediated B cell depletion revealed by intravital imaging. J Clin Invest 123, 5098–5103. 10.1172/JCI70972.

32. Singh, G., Garcia-Bernalt Diego, J., Warang, P., Park, S.C., Chang, L.A., Noureddine, M., Laghlali, G., Bykov, Y., Prellberg, M., Yan, V., et al. (2024). Outcome of SARS-CoV-2 reinfection depends on genetic background in female mice. Nat Commun 15, 10178. 10.1038/s41467-024-54334-7.

33. Umair, M., Fazazi, M.R., and Rangachari, M. (2022). Biological Sex As a Critical Variable in CD4(+) Effector T Cell Function in Preclinical Models of Multiple Sclerosis. Antioxid Redox Signal 37, 135–149. 10.1089/ars.2021.0202.

34. Hirsch, R., Bluestone, J.A., DeNenno, L., and Gress, R.E. (1990). Anti-CD3 F(ab’)2 fragments are immunosuppressive in vivo without evoking either the strong humoral response or morbidity associated with whole mAb. Transplantation 49, 1117–1123. 10.1097/00007890-199006000-00018.

35. Youness, A., Miquel, C.H., and Guery, J.C. (2021). Escape from X Chromosome Inactivation and the Female Predominance in Autoimmune Diseases. Int J Mol Sci 22. 10.3390/ijms22031114.

36. Hoffmann, J.P., Liu, J.A., Seddu, K., and Klein, S.L. (2023). Sex hormone signaling and regulation of immune function. Immunity 56, 2472–2491. 10.1016/j.immuni.2023.10.008.

37. Chi, L., Liu, C., Gribonika, I., Gschwend, J., Corral, D., Han, S.J., Lim, A.I., Rivera, C.A., Link, V.M., Wells, A.C., et al. (2024). Sexual dimorphism in skin immunity is mediated by an androgen-ILC2-dendritic cell axis. Science 384, eadk6200. 10.1126/science.adk6200.

38. Wang, X., Liu, Z., Lu, L., and Jiang, S. (2022). The putative mechanism of lymphopenia in COVID-19 patients. J Mol Cell Biol 14. 10.1093/jmcb/mjac034.

39. O’Brien, B.A., Huang, Y., Geng, X., Dutz, J.P., and Finegood, D.T. (2002). Phagocytosis of apoptotic cells by macrophages from NOD mice is reduced. Diabetes 51, 2481–2488. 10.2337/diabetes.51.8.2481.

40. Jun, H.S., Yoon, C.S., Zbytnuik, L., van Rooijen, N., and Yoon, J.W. (1999). The role of macrophages in T cell-mediated autoimmune diabetes in nonobese diabetic mice. J Exp Med 189, 347–358. 10.1084/jem.189.2.347.

41. Lee, M.S., Kwon, H.J., and Kim, H.S. (2012). Macrophages from nonobese diabetic mouse have a selective defect in IFN-gamma but not IFN-alpha/beta receptor pathway. J Clin Immunol 32, 753–761. 10.1007/s10875-012-9682-3.

42. Robertson, S.J., Bedard, O., McNally, K.L., Shaia, C., Clancy, C.S., Lewis, M., Broeckel, R.M., Chiramel, A.I., Shannon, J.G., Sturdevant, G.L., et al. (2023). Genetically diverse mouse models of SARS-CoV-2 infection reproduce clinical variation in type I interferon and cytokine responses in COVID-19. Nat Commun 14, 4481. 10.1038/s41467-023-40076-5.

43. Marquez, E.J., Chung, C.H., Marches, R., Rossi, R.J., Nehar-Belaid, D., Eroglu, A., Mellert, D.J., Kuchel, G.A., Banchereau, J., and Ucar, D. (2020). Sexual-dimorphism in human immune system aging. Nat Commun 11, 751. 10.1038/s41467-020-14396-9.

44. Ucar, D., Marquez, E.J., Chung, C.H., Marches, R., Rossi, R.J., Uyar, A., Wu, T.C., George, J., Stitzel, M.L., Palucka, A.K., et al. (2017). The chromatin accessibility signature of human immune aging stems from CD8(+) T cells. J Exp Med 214, 3123–3144. 10.1084/jem.20170416.

45. Penaranda, C., Tang, Q., and Bluestone, J.A. (2011). Anti-CD3 therapy promotes tolerance by selectively depleting pathogenic cells while preserving regulatory T cells. J Immunol 187, 2015–2022. 10.4049/jimmunol.1100713.

46. Tian, J., Azhir, A., Decaro, M., Chau, N., Hugel, J., Morris, M., Cheng, J., Fard, P., Bassett, I.V., Bell, D.S., et al. (2026). Long COVID Persistence and Surveillance Gaps Across 58 US Hospitals. JAMA Netw Open 9, e2614909. 10.1001/jamanetworkopen.2026.14909.

47. Narasimhan, H., Cheon, I.S., Qian, W., Hu, S.S., Parimon, T., Li, C., Goplen, N., Wu, Y., Wei, X., Son, Y.M., et al. (2024). An aberrant immune-epithelial progenitor niche drives viral lung sequelae. Nature 634, 961–969. 10.1038/s41586-024-07926-8.

48. Ma, T., Ryu, H., Yin, K., Dalhuisen, T., Robbins, D., Bailey, T.T., Thomas, S.A., Fehrman, E.A., Kelly, J.D., Martin, J.N., et al. (2026). Persistent cytolytic CD8(+) T cells recognize SARS-CoV-2 and herpesvirus epitopes in long COVID. Cell Rep Med 7, 102946. 10.1016/j.xcrm.2026.102946.

49. Waltz, M., Lyerly, A.D., and Fisher, J.A. (2023). Exclusion of Women from Phase I Trials: Perspectives from Investigators and Research Oversight Officials. Ethics Hum Res 45, 19–30. 10.1002/eahr.500170.

50. Zucker, I., and Prendergast, B.J. (2020). Sex differences in pharmacokinetics predict adverse drug reactions in women. Biol Sex Differ 11, 32. 10.1186/s13293-020-00308-5.

