## Supplementary Materials for "Male and female mice achieve equivalent protection against heterologous SARS-CoV-2 infection through divergent immune mechanisms"

### FOR

**CONTENT**

**1. Supplementary Figures (4 Figures)**

**Figure S1.** Protection from heterologous Delta infection is reliant on active infection and is antigen-specific.

**Figure S2.** Neutralizing antibodies and circulating B cells are not sufficient for protection from lethal secondary Delta infection.

**Figure S3.** Opt-SNE analysis and clustering of cell populations from flow cytometry.

**Figure S4.** Female protective responses to heterologous infection are distinct from males.

**2. Supplementary Materials and Methods**

**3. Supplementary References**

#### SUPPLEMENTARY FIGURES

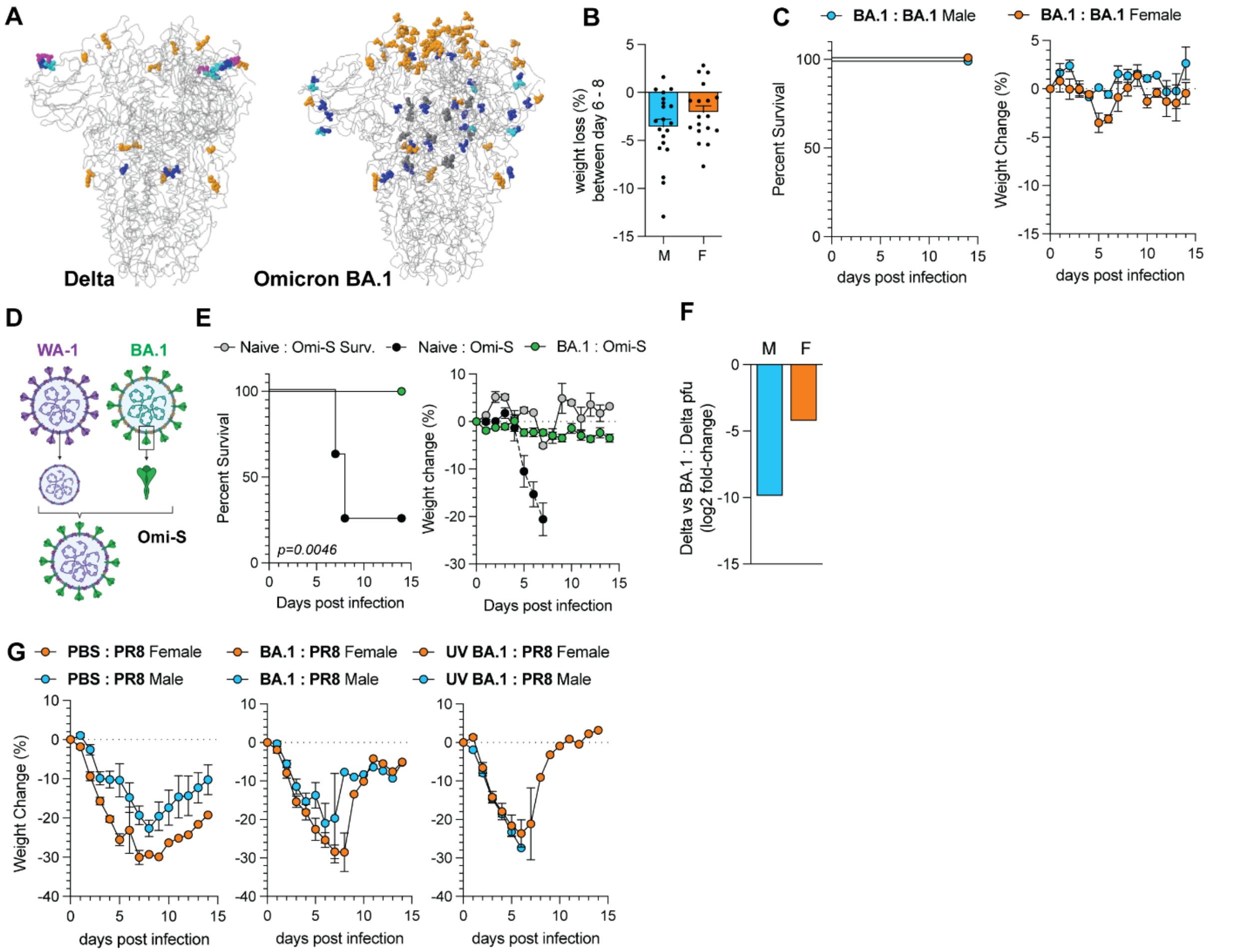


**Figure S1: Protection from heterologous Delta infection is reliant on active infection and is antigen-specific. (A)** Structure of SARS-CoV-2 spike protein from Delta and Omicron BA.1 showing mutations relative to the ancestral strain. Structure and analysis and spike structural visualization were performed using *nextsrain.org.* **(B)** Bar plot showing average weight change between days 6-8 in female (n=15) and male (n=17) mice that were challenged with $10^4$ pfu of Omicron BA.1. Ordinary One-Way ANOVA. Error bars indicate S.E.M. **(C)** Survival and weight loss of BA.1-exposed male (n=3) and female (n=3) mice after BA.1 secondary challenge. Mantel-Cox log-rank test. **(D)** Schematic of Omi-S virus depicting Wuhan-1 virus harboring Omicron BA.1 spike. **(E)** Survival and weight loss of BA.1exposed male (n=7) and female (n=8) mice after Omi-S secondary challenge. Mantel-Cox log-rank test. **(F)** $Log_2$ fold change between mean viral titer in the lungs of naïve and BA.1-exposed mice 2 days post Delta secondary infection. **(G)** Weight loss of mock (left; n: m= 7, f=8), BA.1-exposed (middle; n: m= 6, f=7), or UV-BA.1 exposed (right; n: m= 6, f=7) male and female mice after secondary infection with $1x10^3$ pfu of Influenza PR8 (H1N1). Mantel-Cox log-rank test.

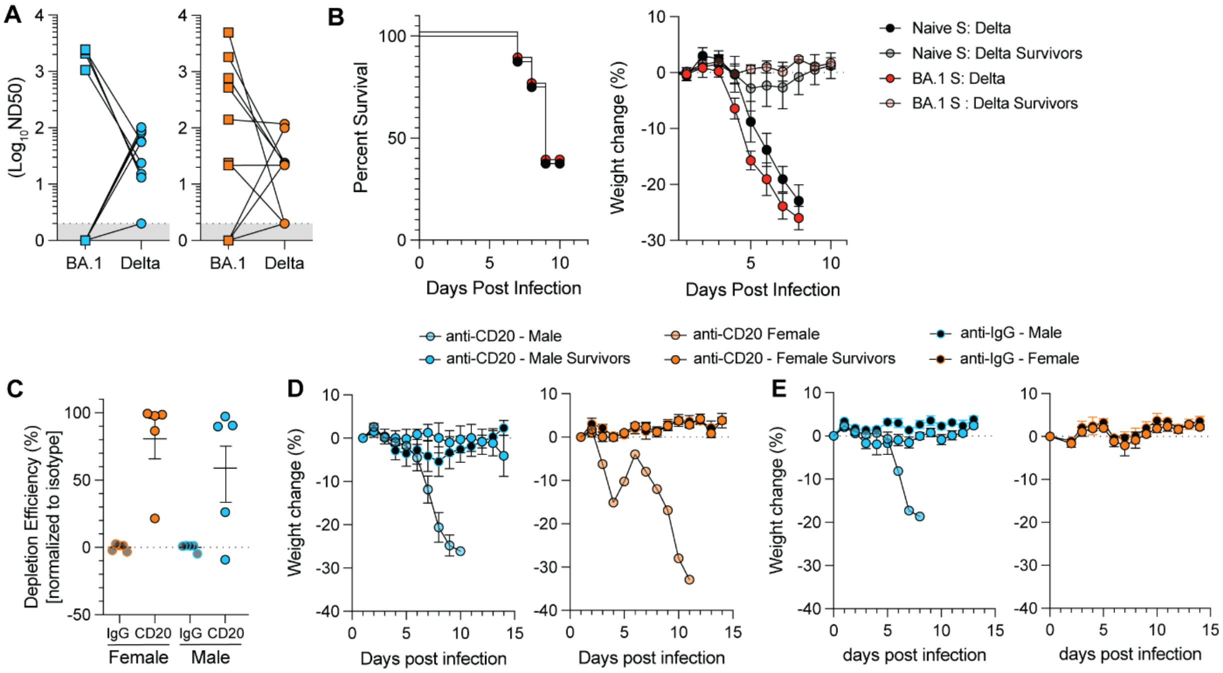


**Figure S2: Neutralizing antibodies and circulating B cells are not sufficient for protection from lethal secondary Delta infection. (A)** Correlation of BA.1 and Delta neutralizing antibodies in male (left) and female (right) mice at 30 days post BA.1 primary infection. Connecting lines indicate NAbs of BA.1 and Delta from the same mouse. **(B)** Survival (left) and weight change (right) after Delta infection of mice that received intravenous administration of 150 uL of serum from BA.1 (n = 8) or mock mice (n = 8) 6 hours prior to infection. Survival analysis was performed by the Mantel-Cox log-rank test. Weight loss analysis was performed by multiple t-tests with Bonferroni correction for multiple comparisons. Error bars indicate S.E.M. **(C)** Depletion efficiency in blood of male or female mice treated with anti-CD20 or anti-IgG. Analysis was performed on peripheral mononuclear cells from the blood of mice and analyzed by flow cytometry. Multiple t-tests with Bonferroni correction for multiple comparisons. Error bars indicate S.E.M. **(D)** Weight loss after Delta challenge of mice depleted with anti-CD20 (n: m=5, f=8) or anti-IgG (n: m=5, f=7) 1 day prior to BA.1 primary infection. Multiple t-tests with Bonferroni correction for multiple comparisons. Error bars indicate S.E.M. **(E)** Weight loss after Delta challenge of mice depleted with anti-CD20 (n: m=5, f=5) or anti-IgG (n: m=5, f=5) 24 hours prior to Delta secondary infection. Multiple t-tests with Bonferroni correction for multiple comparisons. Error bars indicate S.E.M.

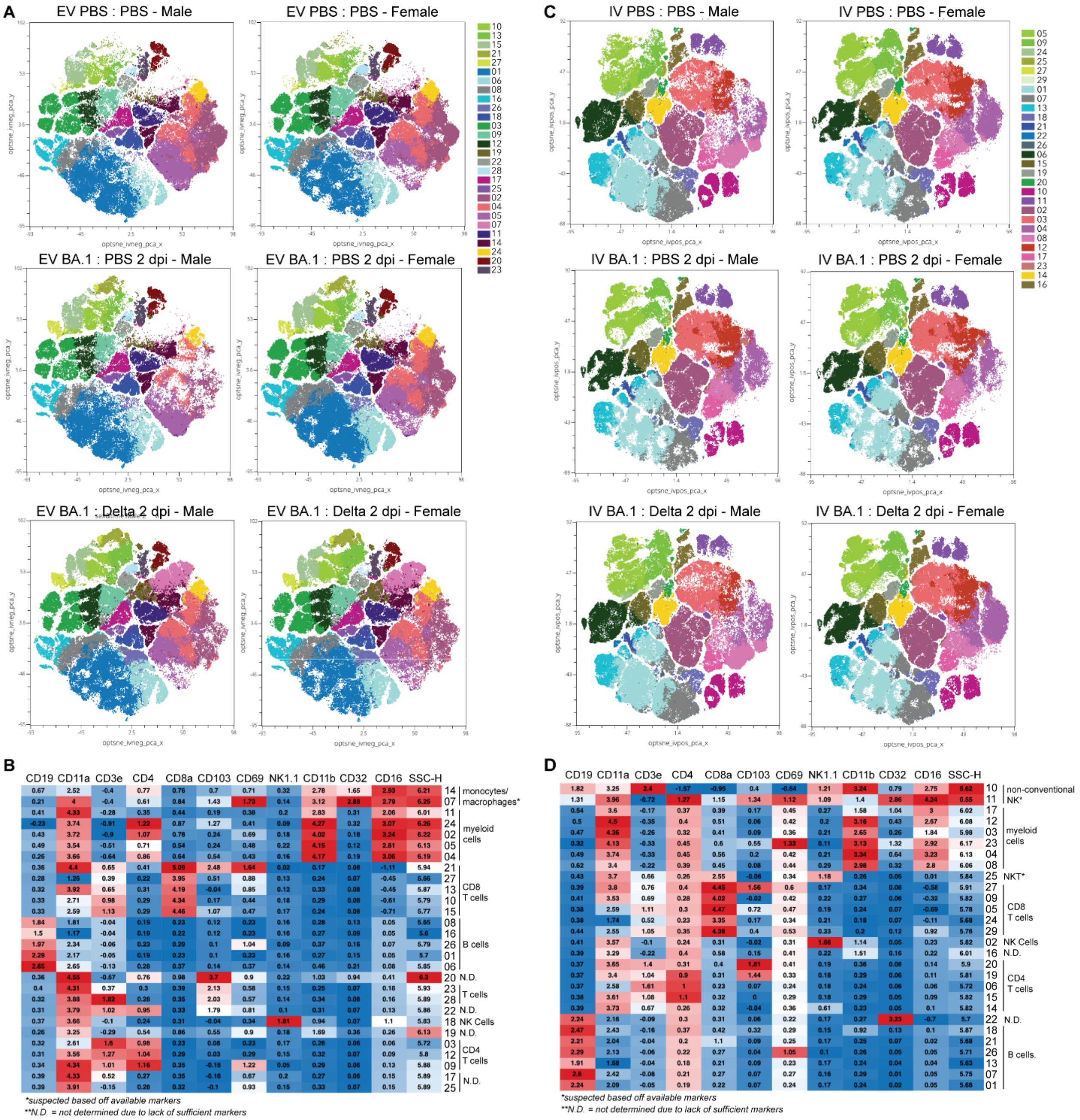


**Figure S3: Opt-SNE analysis and clustering of cell populations from flow cytometry. (A-D)** Flow cytometry analysis was performed on the lungs of male and female mice 32 days post BA.1 challenge, 2 days post Delta challenge in BA.1-exposed mice, or mock-infected mice. **(A)** Lung extravascular (i.v. CD45.2$^{neg}$CD45$^{pos}$) immune compartment of mock, 32-days post BA.1 challenge (BA.1 32 dpi), or 2 days post Delta challenge of BA.1-exposed (BA.1 : Delta 2 dpi) mice. Spectral cytometry data projected into opt-SNE space and clustered with

Phenograph. **(B)** Heatmap showing normalized expression of each marker in Phenograph clusters for the lung extravascular compartment. Markers are listed above the columns, and cluster names are on the right Y-axis. **(C)** Lung intravascular (i.v. CD45.2$^{pos}$CD45$^{neg}$) immune compartment of mock, 32-days post BA.1 challenge (BA.1 32 dpi), or 2 days post Delta challenge of BA.1-exposed (BA.1 : Delta 2 dpi) mice. Spectral cytometry data projected into opt-SNE space and clustered with Phenograph. **(D)** Heatmap showing normalized expression of each marker in Phenograph clusters for the lung intravascular compartment. Markers are listed above the columns, and cluster names are on the right Y-axis.

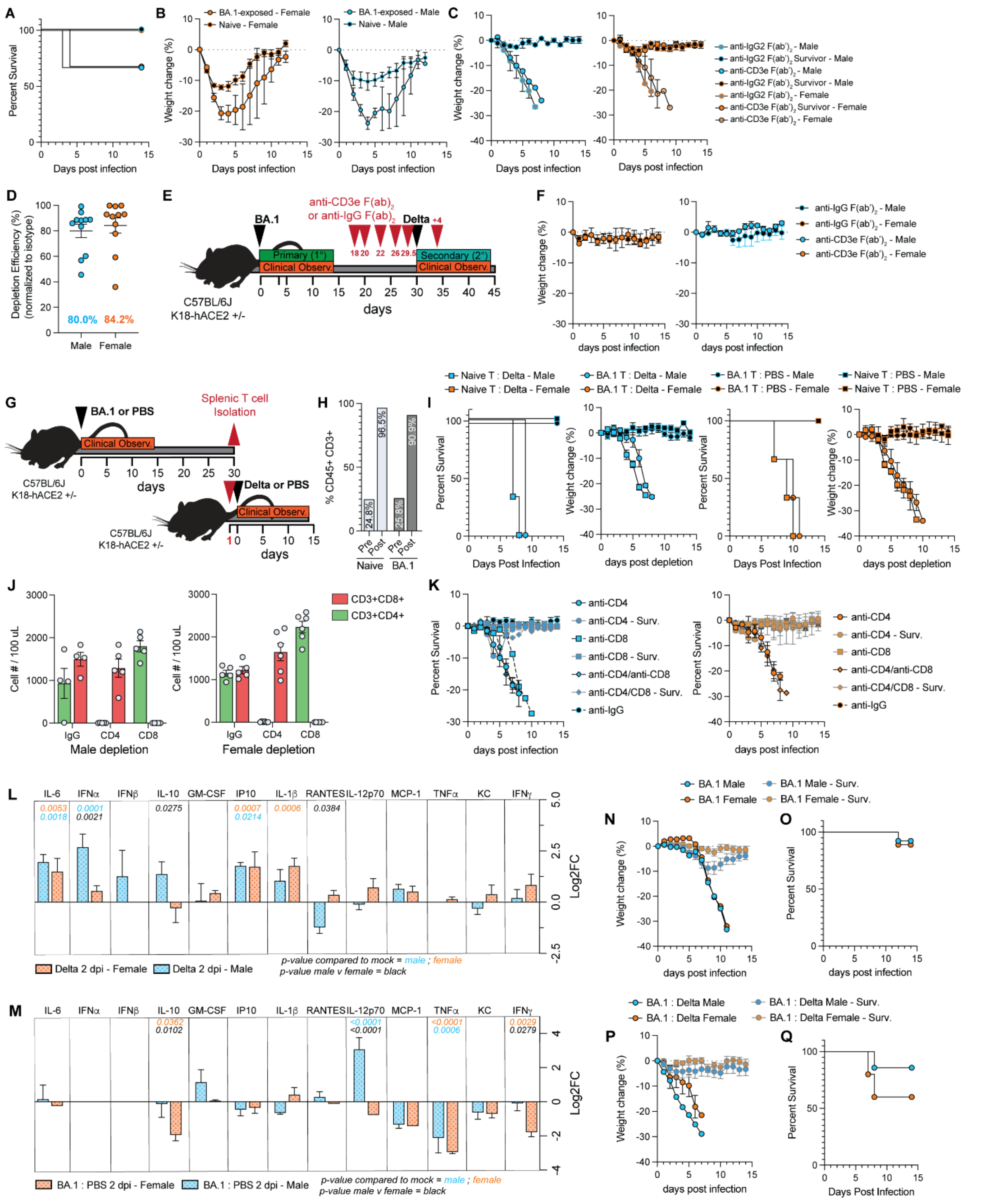


**Figure S4: Female protective responses to heterologous infection are distinct from males.** **(A)** Survival curve and **(B)** weight loss of BA.1-exposed and naïve mice treated with anti-CD3e (n: m=3, f=3) or anti-IgG (n: m=3, f=3). Survival analysis was performed by the Mantel-Cox log-rank test. Weight loss analysis was performed by multiple t-tests with Bonferroni correction for multiple comparisons. Error bars indicate S.E.M. **(C)** Weight loss for Delta secondary infection of mice depleted with anti-CD3e F(ab')$_2$ (n: m=7, f=7) or anti-IgG F(ab')$_2$ (n: m=7, f=7) prior to and during BA.1 primary infection. Multiple t-tests with Bonferroni correction for multiple comparisons. Error bars indicate S.E.M. **(D)** Depletion efficiency in blood of male or female mice treated with anti-CD3e F(ab')$_2$ or anti-IgG F(ab')$_2$. Analysis was performed on peripheral mononuclear cells from the blood of mice and analyzed by flow cytometry. Ordinary One-Way ANOVA. Error bars indicate S.E.M. **(E)** Schematic for depletion using anti- CD3e F(ab')$_2$ or anti-IgG F(ab')$_2$ after BA.1 primary infection. **(F)** Weight loss after Delta challenge of mice depleted with anti-CD3e F(ab')$_2$ (n: m=3, f=3) or anti-IgG F(ab')$_2$ (n: m=3, f=3) after BA.1 primary infection and during Delta secondary infection. Multiple t-tests with Bonferroni correction for multiple comparisons. Error bars indicate S.E.M. **(G)** Schematic for splenic T cell transfer experiments. $1x10^6$ T cells were injected intraperitoneally. **(H)** Single cells were generated from the spleen, and T cells were purified using a Pan T cell Isolation Kit II (negative selection) from Miltenyi Biotec. Percentage of CD45+CD3+ cells pre-purification and post-purification. Enriched T cells post-purification were used for adoptive transfer experiments. **(I)** Survival and weight loss upon Delta or mock challenge of male (blue) and female (orange) mice that were administered BA.1-exposed or naïve splenic T cells one day prior to challenge. (n: m=3 f=3 for all conditions). Survival analysis was performed by Mantel-Cox log-rank test. Weight loss analysis was performed by multiple t-tests with Bonferroni correction for multiple comparisons. Error bars indicate S.E.M. **(J)** Depletion efficiency in blood of male or female mice treated with anti-CD3e F(ab')$_2$ or anti-IgG F(ab')$_2$. Peripheral mononuclear cells from mouse blood were analyzed by flow cytometry. **(K)** Weight loss after Delta challenge of mice depleted with anti-CD4 (n: f=10, f=7), anti-CD8 (n: m=8, f=9), anti-CD4/anti-CD8 (n: m=4, f=5), or anti-IgG F(ab')$_2$ (n: m=7, f=8) after BA.1 primary infection and during Delta secondary infection. Multiple t-tests with Bonferroni correction for multiple comparisons. Error bars indicate S.E.M. (**L-M**) BioLegend LegendPlex 13-plex antiviral cytokine quantification from serum collected at (**L**) 2 days post primary Delta challenge (n: m=2, f=5). or (**M)** 30 days post BA.1 challenge (n: m=3, f=3). $Log_2$Fold-change normalized to serum from mock. Ordinary Two-way ANOVA with Šídák correction for multiple comparisons. Error bars indicate S.E.M. (**N-Q**) 85–95-week-old male and female mice were challenged with BA.1 30 days prior to secondary challenge with Delta. (**N**) Weight loss and (**O**) survival after BA.1 primary challenge (n: m= 13, f= 9). (**P**) Weight loss and (**Q**) survival after Delta secondary challenge (n: m= 7, f= 5). Survival analysis was performed by the Mantel-Cox log-rank test. Weight loss analysis was performed by multiple t-tests with Bonferroni correction for multiple comparisons. Error bars indicate S.E.M.

#### SUPPLEMENTARY MATERIALS AND METHODS

**Cell lines.** VeroE6 cells were grown in Dulbecco's modified Eagle's medium (DMEM) supplemented with 10% heat-inactivated fetal bovine serum (FBS) (Bio-Techne, R&D Systems, Minneapolis, MN, USA) and 1% (v/v) Penicillin-Streptomycin (P/S) (Thermo Scientific, Waltham, MA, USA). Caco-2 and A549 cells expressing human ACE2 and human TMPRSS2 (Caco-2 and A549 hACE2/hTMPRSS2) were previously described[1]. These cells were both maintained in DMEM containing 10% FBS and 1% P/S supplemented with 2.5 ug/mL of Puromycin and 2.5 µg/mL of Blasticidin. Madin-Darby Canine Kidney (MDCK) cells were maintained in Minimum Essential Medium supplemented with 10% FBS and 1% P/S.

**SARS-CoV-2 Virus Stocks.** SARS-CoV-2 B.1.617.2 (Delta) and Omicron BA.1 clinical isolates were kindly provided by the laboratory of Dr. John H. Connor at Boston University. SARS-CoV-2 Omi-S was generated using a circular polymerase extension reaction (CPER), as previously described[2]. SARS-CoV-2 Delta virus was grown and titrated on VeroE6 cells. Omicron BA.1 and Omi-S recombinant viruses were grown and titrated on Caco2 hACE2/hTMPRSS2 cells. For growth, $8x10^6$ cells were plated into a T-175 flask the day prior to infection. The next day, cells were infected with a multiplicity of infection (moi) = 0.1 pfu/cell of passage 1 or 2 (P1 or P2) Delta, P2 BA.1, or P1 Omi-S in 10 mL of Opti-MEM media + GlutaMax (Gibco®, Carlsbad, CA, USA) and incubated for 1 hr at 37°C to allow for viral adsorption. After 1 hr incubation, 10 mL of DMEM with 4% FBS was added to cells, incubated for 24 hr, and then the media was replaced with 25 mL of DMEM with 2% FBS and incubated at 37°C. Cells were assessed for cytopathic effect (CPE) and virus was collected when significant CPE was observed (between 60-96 hr post infection).

Cell supernatant was clarified by centrifugation for 10 min at 350 *x g* at 4°C and filtered through a 0.22 µM filter. Delta virus was overlaid on a 20% sucrose cushion (Sigma-Aldrich, St. Louis, MO, USA) and concentrated by ultracentrifugation (Beckman Coulter Optima L-100k; SW32 Ti rotor) at 25,000 RPM for 2 hours at 4°C. Viral pellets were then suspended in 1x PBS and titrated by plaque assay on Vero E6 cells (see *Viral quantification by plaque assay*). For Omicron BA.1 and Omi-S, virus was mixed at a 1:1 ratio with PEG6000 (Sigma-Aldrich, cat# 1546580) and incubated at 4°C overnight, centrifuged at 10,000 RPM for 30 min at 4°C, then suspended in 1xPBS.

**Influenza PR8 viral stocks.** A/Puerto Rico/8/1934 (H1N1) (PR8) virus was generated in a BSL-2 from a P1 virus stock. For growth, $8x10^6$ MDCK cells were plated into a T-175 flask the day prior to infection. The next day, media was removed, cells were washed with 1xPBS twice, and then infected with a multiplicity of infection (MOI) = 0.1 pfu/cell of Influenza PR8 (H1N1) in 10 mL Opti-MEM + Glutamax. After 1 hr incubation at 37°C to allow for viral adsorption, the medium was removed, and 25 mL of 1xMEM (Gibco, cat# 11095080) without FBS was added. Cells were monitored for CPE and collected when significant CPE was observed. For purification, cell supernatant was clarified by centrifugation for 10 min at 350 *x g* at 4°C and filtered through a 0.22 uM filter. Virus was overlaid on a 20% sucrose cushion (Sigma-Aldrich, St. Louis, MO, USA) and concentrated by ultracentrifugation (Beckman Coulter Optima L-100k; SW32 Ti rotor) at 25,000

RPM for 2 hours at 4°C. Viral pellets were then suspended in 1x PBS titrated by plaque assay on MDCK cells.

**UV inactivation of SARS-CoV-2 BA.1.** Prior to inoculation, 1 mL of prepared inoculation SARS-CoV-2 BA.1 mix (1x$10^4$ pfu per mouse in 50 µL) was placed into a well of a 12-well plate and placed into a Stratagene UV Stratalinker 2400 (Stratagene [now Agilent Technologies], cat# 400075) and exposed twice to 2300 Joules. Inactivation of the virus was confirmed by plaque assay analysis.

**Mouse strains.** 15–27-week-old male and female heterozygous K18-hACE2 C57BL/6J (2B6.Cg-Tg(K18-ACE2)2Prlmn/J; Jackson Laboratory, Catalog # 034860) and 12-24-week-old male and female heterozygous NSG-hACE2 (NOD.Cg-Tg(K18-ACE2)2Prlmn $Prkdc^{scid}Il2rg^{tm1Wjl}$/J; Jackson Laboratory, Catalog # 034901) were used in these studies. For aged studies 85-95-week-old male and female heterozygous K18-hACE2 C57BL/6J mice were used. Animals were group-housed by sex in Tecniplast green line individually ventilated cages (Tecniplast, Buguggiate, Italy). Mice were maintained on a 12:12 light cycle at 30–70% humidity and provided *ad libitum* water and standard chow diets (LabDiet, St. Louis, MO, USA).

**Clinical monitoring and euthanasia criteria.** For survival studies, mice were monitored for changes in body weight, altered respiration, general changes in appearance, responsiveness and neurological signs of disease. An IACUC-approved clinical-scoring system was used to monitor disease progression and define humane endpoints. The score of "1" was given for each of the following situations: body weight, 10–29% loss; respiration, rapid and shallow with increased effort; appearance, ruffled fur and/or hunched posture; responsiveness, low to moderate unresponsiveness; and neurological signs, tremors. The sum of these individual scores constituted the final clinical score (0-5). Mice were humanely euthanized in case of weight loss greater than or equal to 30%, or if they received a clinical score of 4 or above for two consecutive days or were considered moribund. Body weight and clinical score were recorded once daily throughout the study. For survival curves, mice euthanized on a given day were counted as dead the day after. Mice that were found dead in the cage were counted dead on the same day. For euthanasia, an overdose of ketamine was administered followed by a secondary method of euthanasia.

**Tissue collection.** At predetermined endpoints for tissue collection, mice were euthanized by ketamine/xylazine overdose. Tissues were then collected and placed in 600-750 µL of RNAlater (MilliporeSigma: #R0901500ML) and stored at -80C for downstream analysis. Blood was collected by cardiac puncture after euthanasia and placed into a 1.5 mL Eppendorf tube. After collection, blood was allowed to clot for 20-30 minutes at room temperature, then centrifuged for 15 minutes at 3500 RPM in a benchtop centrifuge. Serum was then collected, transferred to a new tube, and stored at -80C prior to downstream analysis or use.

***In vivo* antibody-mediated cell depletions.** All depletions were performed in 15-20-week-old male and female K18-hACE2 mice.

*T cell depletion*: For depletions prior to primary infection, mice were administered 200 ug of inVivoMAb anti-CD3e F(ab’)$_2$ (clone 145-2C11 f(ab')2 Fragments; BioXcell cat# BE0001-1FAB) intraperitoneally at 1, 4, 8, 10, and 12 days prior and 4 days post primary SARS-CoV-2 BA.1 infection. Isotype anti-Armenian hamster inVivoMAb hamster IgG f(ab')2 fragment (BioXcell cat # BE0091-FAB). For T cell depletions post primary infection, mice were administered 200 ug of inVivoMAb anti-CD3e F(ab’)$_2$ intraperitoneally at 1, 4, 8, 10, and 12 days prior to secondary infection. Isotype inVivoMAb hamster IgG f(ab')2 fragment for all T cell depletion experiments.

*B cell depletions*: Mice were injected intraperitoneally with 200 ug of inVivoMAb anti-CD20 (clone MB20-11; BioXcell cat# BE0356) 24-hours prior to primary SARS-CoV-2 BA.1 infection or secondary SARS-CoV-2 Delta infection. An InVivoMAb mouse IgG2c, κ isotype control, anti-dengue virus (clone DV5-1; BioXcell cat#BE0366) was used as a control for B cell depletion experiments.

*CD4 and CD8 depletions:* Mice were injected intraperitoneally with 200 ug of inVivoMAb anti-CD4 (clone GK1.5; BioXcell Cat# BE0003-1), inVivoMAb anti-CD8 (clone 2.43; BioXcell Cat# BE0061), 200 ug each of inVivoMAb anti-CD4 and inVivoMAb anti-CD8 (co-depletion), or inVivoMAb anti-IgG2b κ (clone LTF-2; BioXcell Cat# BE0090) at 1, 3, and 5 days prior to BA.1 infection and 4 days post-BA.1 infection.

*Validation of depletion:* Blood was collected via submandibular bleeding or heart puncture and transferred into EDTA capillary collection tubes (Microvette 600 K3E; Sarstedt, Nümbrecht, Germany). 100 µL of blood was centrifuged at 3,500 RPM for 10 minutes at room temperature; serum was removed, and whole blood was subjected to ACK lysis for 10 minutes at room temperature. Lysis was quenched with RPMI media with 2% FBS, and cells were washed twice with 1x PBS with 1% FBS solution before downstream processing. After lysis, cells were centrifuged at 300 X g for 8 min at 4°C. The cell pellet was resuspended in 50 µL Zombie NearIR (1:800 dilution) live/dead staining in 1x PBS and incubated for 20 minutes on ice. Cells were then washed twice with 200 µL of 1xPBS with 1% FBS. After washing, cells were incubated with 25 µL of BD stain buffer with 1 µL of TruStain FcX (anti-mouse CD16/32 antibody) (Biolegend, cat# 101320) and 2.5 µL True-Monocyte blocking buffer (Biolegend, cat# 426103) and incubated for 10 min at room temperature. After blocking, 25 µL of antibody cocktail containing (anti-CD4 PerCP, anti-CD8 PE-Cy7, anti-CD19 BV605, anti-CD45.2 FITC, and anti-CD3e Pacific Blue (clone 17A2, Biolegend, cat#100214, 1:100)) in 1xPBS with 1% FBS was added to each sample and incubated in the dark for 30 min at 4°C. After incubation, cells were washed twice in 1xPBS with 1% FBS and analyzed by flow cytometry. Flow cytometric analysis for depletion validation was performed using an LSRII Flow Cytometer (BD Biosciences) and analyzed using FlowJo software v10 (TreeStar, Ashland, OR, USA).

***In vivo* intranasal IFNγ treatments.** 12–22-week-old NSG-K18-hACE2 mice were intranasally infected with $1x10^4$ pfu of SARS-CoV-2 Delta virus prior to treatment. For treatment, mice were put under anesthesia with 1-3% isoflurane and intranasally administered 2 µg of IFNγ (R&D Systems; Cat # 485-MI) in 50 µL of PBS or PBS alone. For early treatment, administration of IFNγ began at 1 day post-infection and was given daily until 7 days post-infection. For late treatments, IFNγ treatment was started at 4 days post-infection and continued until 7 days post-infection. Mice in late treatment groups were treated with PBS on days 1, 2, and 3 post-infection. Note that the use of daily isoflurane anesthesia can affect survival during infection but is required for intranasal treatments.

**Viral quantification by plaque assay.** *SARS-CoV-2:* The titer of our viral stocks was determined by plaque assay. For Delta stock, titers were determined on Vero E6 cells. For Omicron and Omi-S stocks, titers were determined on Caco-2 cells overexpressing hACE2/hTMPRSS2 (Caco2 A/T). For VeroE6 cells, cells were seeded into a 12-well plate at a density of $2x10^5$ cells per well. For Caco-2 A/T cells, cells were seeded into a 12-well plate at a density of $2x10^5$ cells per well. The next day, cells were infected with 10-fold serial dilutions of the viral stocks and incubated for 1 hour at 37°C to allow for viral adsorption. After 1 hour of incubation, 1 mL of overlay media (1.2% Avicel (DuPont, Wilmington, DE, USA; RC-581) in DMEM supplemented with 2% FBS and 1% Pen/Strep) was added to each well. Three days later, the overlay medium was removed, and cells were fixed for 1 hour at room temperature in 10% neutral-buffered formalin (ThermoFisher Scientific). Formalin was removed after fixation and cells were stained for 30 min at room temperature with 0.1% crystal violet (Sigma-Aldrich) prepared in 10% ethanol/water. Crystal violet was removed; cells were washed with water, and plaque-forming units were counted to determine viral titers.

*Influenza PR8*: MDCK cells were used for plaque-forming assays to measure the PR8 viral titer. PR8 stock was serially diluted 1:10 in Opti-MEM, then each dilution was plated onto 12-well plates and incubated at 37 °C with 5% CO2 for 1 hour, followed by the addition of 1 mL of 1.2% Avicel mixed with 1x MEM (without FBS) containing 1 μg/mL of trypsin-TPCK (L-(tosylamido-2-phenyl) ethyl chloromethyl ketone) (Worthington Biochemicals, Lakewood, NJ, USA). Cells were incubated for three days before the overlay was removed. Cells were then fixed with 10% neutral buffered formalin for 1 hour at room temperature and stained with crystal violet (0.1% crystal violet in 10% ethanol/water).

**Tissue processing for viral quantification.** 40-70 mg of tissue was weighed out and placed into a 2 mL tube with a 5 mm stainless steel bead and 600 μL of OptiMEM. Tissues were homogenized using a Qiagen TissueLyser II (Qiagen) with the following cycle: two min dissociation at 1800 oscillations/min, one min rest, two min dissociation at 1800 oscillations/min. Dissociated tissues were centrifuged at 13,000 rpm for 10 min at room temperature, and the cleared supernatant was transferred to a new 1.5 mL Eppendorf tube for downstream analysis. Lung titers were determined by the following formula:

$$\frac{\boldsymbol{pfu}}{\boldsymbol{mg}} = \frac{\left(\frac{\boldsymbol{\#\ of\ plaques}}{\boldsymbol{dilution * plating\ volume\ (mL)}}\right) * \mathbf{0.600}\ \boldsymbol{mL}}{\boldsymbol{mg\ of\ tissue}}$$

**Serum collection.** Blood was collected by cardiac puncture after euthanasia, placed in 1.5 mL Eppendorf tubes, and allowed to clot at room temperature for 20-30 minutes. Blood was centrifuged at 3500 RPM for 10 minutes in a benchtop centrifuge. Serum was removed from the top layer, transferred to a new 1.5 mL Eppendorf tube, and stored at -80°C.

**Serum neutralization assay.** One day prior to the experiment, $2x10^4$ A549-hACE2/hTMPRSS2 cells were plated into a 96-well plate. Serum was decomplemented at 56 °C for 30 min and an initial dilution of 1:10 was prepared in OptiMEM. Two-fold dilutions were subsequently prepared and mixed with SARS-CoV-2 Delta or Omicron BA.1 virus (MOI = 1) for 1 hr at room temperature before being plated onto cells. After a 1-hr incubation at 37 °C, the inoculum was removed, and 200 µL of fresh DMEM containing 2% FBS and 1% penicillin/streptomycin was added. After a 36 hr incubation at 37 °C with 5% CO2 media was removed and cells were fixed with 10% formalin for 1 hr. Cells were then washed three times with 1xPBS and permeabilized for 30 minutes with 1xPBS containing 1% Triton-X100 at room temperature. After permeabilization, cells were blocked with 1xPBS containing 0.1% Triton-X100 and 5% BSA for 30 minutes at room temperature. After blocking, cells were incubated with anti-SARS-CoV-2 nucleocapsid (N) antibody (Rockland Antibodies, cat# 200-401-MS4) (1:2000) in 1xPBS with 0.1% Triton-X and 1% BSA overnight at 4°C. Antibody was then removed and washed three times with 1xPBS prior to secondary antibody staining with goat anti-Rabbit IgG (H+L) Cross-Adsorbed Secondary Antibody, Alexa Fluor 568 (1:1000) (ThermoFisher, cat# A-11011) for 1 hr at room temperature, protected from light. Cells were then washed three times with 1x PBS and counterstained with Hoechst. Quantification of infection was performed by image capture with a Biotek Cytation Imager and analyzed using CellProfiler.

**Serum transfer experiments.** Serum from male and female 15–27-week-old K18-hACE2 mice was collected at 28-30 days post-infection with $1x10^4$ pfu of SARS-CoV-2 BA.1 virus. Serum from both sexes was pooled and used for injection. Transfer experiments were either performed intraperitoneally (I.P.) or intravenously (I.V.). For I.P. transfers, 200 µL of serum was injected 6 hours prior to infection with 1×10^4 pfu of SARS-CoV-2 Delta or Omi-S virus. For I.V. transfers, 150 µL of serum was injected 6 hours prior to infection with 1×10^4 pfu of SARS-CoV-2 Delta virus. For both I.P. and I.V., naïve serum collected from aged and sex matched K18-hACE2 mice was used as a negative control. After infection, mice were monitored for signs of clinical disease and lethality for 14 days or until mice died.

**Preparation of splenic T cells for adoptive transfer.** T cells were isolated from the spleen of male and female K18-hACE2 mice collected at 30 days post-infection with $1x10^4$ pfu of SARS-CoV-2 BA.1 virus. To isolate splenic T cells, spleen was removed and placed into a 70 uM strainer on a 50 mL conical tube. Using a plunger from a 3 mL syringe, the spleen was mashed through the strainer, periodically washing with RPMI media (Gibco; cat# 11875093) containing 2% FBS. Once through the strainer, cells were pelleted by centrifugation at 350 *x g* for 8 minutes at 4°C, washed with 5 mL of 1x PBS, then red blood cells were lysed using 3 mL of ACK lysis buffer (Gibco; cat# A1049201) for 10 minutes at room temperature. Lysis was neutralized using 22 mL RPMI 2% FBS prior to centrifugation at 350 *x g* for 8 minutes at 4°C. Cells were then washed twice with 10 mL of 1xPBS containing 2% FBS and 0.02% Sodium Azide before T cell purification.

**T cell purification.** T cells were purified using a MACS Miltenyi Pan-T cell isolation kit (Miltenyi Biotech, cat# 130-095-130). Purification was performed as described in the manufacturer's protocol using an AutoMACs Pro separator (Miltenyi Biotech, Auburn, CA, USA). Notably, this kit is a negative selection kit that uses a cocktail of biotin-conjugated antibodies against CD11b, CD11c, CD19, CD45R (B220), CD49b (DX5), CD105, Anti-MHC-class II, and Ter-119, thereby preventing T cell activation during purification. T cell purity was assessed by flow cytometry by staining 5% of the purified sample. Purified T cells were counted using a CytoSmart Cell Counter (Corning) prior to transfer.

**T cell transfers.** For transfer experiments, $1x10^6$ purified T cells were injected by intraperitoneal (I.P.) administration into 15-week-old naïve male and female K18-hACE2 mice 24 hours prior to infection. Mice were infected with 1×10^4 pfu of SARS-CoV-2 Delta virus or mock-infected and monitored for 14 days for clinical disease and lethality. As a negative control, T cells were isolated from the spleen

**Single-cell isolation from lungs for flow cytometry.** For flow cytometry analysis, 5-8 minutes prior to euthanasia, mice were retro-orbitally injected with 2ug of anti-CD45.2 BUV737 (clone 104, BD Biosciences, cat # 612778) to allow antibody labeling of circulating immune cells. Then, mice were euthanized, and right lung lobes were collected and stored in RPMI with 2% FBS prior to single-cell suspension generation using a "Lung Dissociation Kit, mouse" (Miltenyi Biotec, Auburn, CA, USA; cat # 130-095-927) per the manufacturer's protocol. Briefly, lungs were placed into gentleMACS™ C Tubes (cat # 130-093-237) containing 2.4 mL of 1X Buffer S, 100 µL of Enzyme D and 15 µL of Enzyme D. Tissues were subject to dissociation using a gentleMACS Dissociator with program m_Lung_01, placed at 37°C for 30 minutes with constant rotation/agitation, then dissociated a second time using a gentleMACS Dissociator with program m_Lung_02. After dissociation, cells were filtered through a 70 µm filter, centrifuged at ~300 x g for 8 minutes at 4°C, supernatant was removed, and cells were suspended in 3 mL of ACK lysis buffer (Gibco, cat # A1049201). Cells were then incubated for 5 minutes at room temperature and lysis of red blood cells was quenched with 10 mL of RPMI with 2% FBS. Cells were then counted and used for downstream flow cytometry staining.

**Staining cells for flow cytometry analysis**. After generation of single-cell suspension, $1x10^6$ cells were used for flow cytometry staining. Cells were centrifuged at 300 X g for 8 min at 4°C. The cell pellet was resuspended in 50 µL Zombie NearIR (BioLegend, cat# 423106, 1:800 dilution) live/dead staining in 1x PBS and incubated for 20 minutes on ice. Cells were then washed twice with 200 µL of BD stain buffer (BD Biosciences, cat# 554656). After washing, cells were incubated with 25 µL of BD stain buffer with 1 µL of TruStain FcX (anti-mouse CD16/32 antibody) (Biolegend, cat# 101320) and 2.5 µL True-Monocyte blocking buffer (Biolegend, cat# 426103) and incubated for 10 min at room temperature. After blocking, 25 µL of antibody cocktail containing antibodies and 10µL BrilliantViolet Staining Buffer (BD Biosciences, cat# 563974) were added to each sample and incubated in the dark for 30 min at 4°C. After staining, 200 µL of BD stain buffer was added to each sample; samples were centrifuged at 300 x g for 8 min at 4°C, washed with 250 µL BD stain buffer, centrifuged at 300 x g for 8 min at 4°C, and then fixed in 200 µL 4% PFA for 1 hour. After fixation, cells were washed twice with 250 µL PBS, resuspended in PBS, and stored protected from light at 4°C until analysis.

**Antibodies for flow cytometry.** Anti-CD45.2 FITC for i.v. (clone: 104, Biolegend, cat #: 109806) 2ug/mouse, anti-NK1.1 BV421 (clone: PK136, Biolegend, cat#: 108731) 1:50, anti-CD45 BV510 (clone: 30-F11, Biolegend, cat#: 103138) 1:200, anti-CD19 BV605 (clone: 6D5, Biolegend, cat#: 115539) 1:200, anti-CD11a BV786 (clone: M17, BD Biosciences, cat# 740866)1:200, anti-CD8 PE-Cy7 (clone: 53-6.7, Biolegend, cat# 100721) 1:800, anti-CD4 PerCP (clone:GK1.5, Biolegend, cat# 100431) 1:50, anti-CD69 PE (clone: H1.2F3, BD Biosciences, cat #: 747481) 1:100, anti-CD11b BUV395 (clone: HL3, BD Biosciences, cat# 563553) 1:100, anti-CD103 APC (clone: 2E7, Biolegend, cat# 367-1031-82) 1:50, anti-CD3 Alexa647 (clone: 145-2C11, Biolegend, cat# 100322) 1:400.

**Flow cytometry Data Acquisition and Analysis.** Flow cytometry analysis was performed using a Cytek Aurora spectral analyzer (Cytek Biosciences). SpectraFlo (Cytek) software was utilized for spectral unmixing of the data using ordinary least square algorithm. Unmixed flow cytometry data were analyzed using OMIQ software from Dotmatics (www.omiq.ai, www.dotmatics.com). Data processing pipeline was assembled within the Omiq.ai cloud computation platform (Dotmatics). Live CD45+CD45.2+ (lung intravascµLar) and live CD45+ CD45.2− (lung extravascµLar) immune cells were concatenated, asinh transformed (cofactor = 6000) and clustered with Phenograph algorithm (k = 30, distance metric = euclidean)[3] and projected into opt-SNE[4] space (perplexity = 30, theta = 0.5, opt-SNE endpoint = 5000; PCA pre-initialization embedding). Clusters were annotated based on phenotypes, color-coded and overlaid on the opt-SNE projection graphs. Each marker's mean fluorescence intensities (MFIs) from clustered datasets were organized into hierarchical heatmaps.

*Cell annotations:* Described cell types were defined by the following: Cells in the intravascular (IV) compartment (Live/Dead-CD45.2+CD45+); extravascular (EV) compartment (Live/Dead-CD45.2-CD45+). Within the EV compartment the following populations were identified: CD4 T cells (CD3+CD19-NK1.1-CD4+), CD4 Tissue resident memory (TRM; CD3+CD19-NK1.1-CD4+CD8-CD11a$^{high}$CD69+), CD4 CD11 low CD69 neg T cells (CD3+CD19-NK1.1-CD4+CD8-CD11a$^{low}$CD69-), CD8 T cells (CD3+CD19-NK1.1-CD4-CD8+), CD8 TRM (CD3+CD19-NK1.1-CD4-CD8+CD103+CD69+), antigen experience CD8 T cells (CD3+CD19-NK1.1-CD4-CD8+CD11a+CD69$^{low}$CD103-), B cells (CD3-CD19+), CD69+ B cells (CD3-CD19+CD69+). Within the IV compartment the following populations were identified: CD4 T cells (CD3+CD19-NK1.1-CD4+CD8-), activated CD4 T cells (CD3+CD19-NK1.1-CD4+CD8-CD69+), CD8 T cells (CD3+CD19-NK1.1-CD4-CD8+), activated CD8 T cells (CD3+CD19-NK1.1-CD4-CD8+CD69+), B cells (CD3-CD19+), CD69+ B cells (CD3-CD19+CD69+).

**Cytokine analysis using BioLegend LegendPlex.** Cytokine analysis was performed on serum that was inactivated using a 1:1 dilution in 2% Tween-80 prepared in 1x PBS for a final concentration of 1% Tween-80. Serum-Tween-80 mixtures were incubated at 4°C for 2 hours to allow for inactivation and then analyzed using a LEGENDplex™ Mouse Anti-Virus Response Panel (13-plex) (Biolegend, cat # 740622) kit as per the manufacturer's protocol. *Of note, while 1% Tween-80 does not abolish the detection of cytokines with this kit, cytokine levels may be reduced due to the presence of Tween-80, though equally for all cytokines.* This kit test for the following cytokines: IFN-γ, CXCL1 (KC), TNF-α, CCL2 (MCP-1), IL-12p70, CCL5 (RANTES), IL-1β, CXCL10 (IP-10), GM-CSF, IL-10, IFN-β, IFN-α, IL-6. Sample collection was performed using a BD Biosciences LSRII Flow cytometer, and data were analyzed using BioLegend Browser-based "Data Analysis Software Suite for LegendPLEX" (https://legendplex.qognit.com/user/login?next=home).

**RNA extraction from tissue.** For RNA extraction, 30–40 mg of tissue was weighed out, placed into a 2 mL tube with 600 µL of RLT buffer with 1% β–mercaptoethanol and a 5 mm stainless steel bead (Qiagen, Hilden, Germany: #69989), then homogenized using a Qiagen TissueLyser II (Qiagen) with the following cycle: two min dissociation at 1800 oscillations/min, one min rest, two min dissociation at 1800 oscillations/min. Homogenized tissues were centrifuged at 13,000 rpm for 10 min at room temperature, and the cleared supernatant was transferred to a new 1.5 mL Eppendorf tube. RNA extractions were performed using a Qiagen RNeasy Plus Mini Kit (Qiagen: #74134), according to the manufacturer's instructions (Qiagen: #79256). RNA was eluted in 30 µL of RNase/DNase free water and quantified by Nanodrop2000.

**Gene expression analysis by RT-qPCR.** RT-qPCR was performed using the Luna Universal One-Step RT-qPCR kit #E3005L, New England Biolabs). The manufacturer's protocol was followed with minor adjustments for a final volume of 12 µL consisting of 2 µL of RNA, 2 µL of forward primer, 2 µL of reverse primer, 0.6 µL of 20X Luna WarmStart RT enzyme mix and 6 µL of 2X Luna universal one-step reaction mix. Using the Applied Biosystems QuantStudio 3 (ThermoFisher Scientific), the 12 µL reaction mixture used the following cycling conditions: 50ºC for 10 minutes (reverse transcription), 95ºC for 2 minutes (initial denaturation) followed by 40 additional cycles of 95ºC for 15 seconds (denaturation) and 60ºC for 1 minute (annealing/extension) and finally a melt curve analysis. The melt curve analysis began with a 30-second waiting period at 65ºC with a temperature change of +0.5ºC/s and a 5-second pause after every increment, during which the machine acquired fluorescence at each new temperature until it reached 95ºC. Threshold cycle values (Ct), also known as quantification cycle (Cq), were determined using QuantStudio Design and Analysis software V1.5.1. Normalization was performed using mouse *Rps11* as a control. *Rps11* primers (Forward: 5'-AAACCGGCAAGGAAAAACTCC-3'; Reverse: 5'-ACCAGTGAAGGGGCATTTCTT-3') *IFNg* primers (Forward: 5'-CAGCAACAGCAAGGCGAAAAAGG-3'; Reverse: 5'-TTTCCGCTTCCTGAGGCTGGA T-3').

**Bulk RNA-sequencing analysis.** Adapters and low-quality bases were removed from raw reads using Trimmomatic (0.39)[5], remaining sequences were mapped to the mouse genome (GRCm39.107) using STAR (2.7.10b)[6] and a count table was produced using subread (2.0.1)[7]. Differentially expressed genes analysis was performed with R (4.1.3)[8] using DESeq2 (1.34.0)[9]. For each comparison, genes with p-val < 0.05 and absolute value of log2 fold change > 1 were considered significant and separated into upregulated and downregulated for overrepresentation analysis using Metascape[10].

#### SUPPLEMENTARY REFERENCES

1. Chen, D.Y., Turcinovic, J., Feng, S., Kenney, D.J., Chin, C.V., Choudhary, M.C., Conway, H.L., Semaan, M., Close, B.J., Tavares, A.H., et al. (2023). Cell culture systems for isolation of

SARS-CoV-2 clinical isolates and generation of recombinant virus. iScience *26*, 106634. 10.1016/j.isci.2023.106634.

2. Chen, D.Y., Chin, C.V., Kenney, D., Tavares, A.H., Khan, N., Conway, H.L., Liu, G., Choudhary, M.C., Gertje, H.P., O'Connell, A.K., et al. (2023). Spike and nsp6 are key determinants of SARS-CoV-2 Omicron BA.1 attenuation. Nature *615*, 143-150. 10.1038/s41586-023-05697-2.
3. Levine, J.H., Simonds, E.F., Bendall, S.C., Davis, K.L., Amir el, A.D., Tadmor, M.D., Litvin, O., Fienberg, H.G., Jager, A., Zunder, E.R., et al. (2015). Data-Driven Phenotypic Dissection of AML Reveals Progenitor-like Cells that Correlate with Prognosis. Cell *162*, 184-197. 10.1016/j.cell.2015.05.047.
4. Belkina, A.C., Ciccolella, C.O., Anno, R., Halpert, R., Spidlen, J., and Snyder-Cappione, J.E. (2019). Automated optimized parameters for T-distributed stochastic neighbor embedding improve visualization and analysis of large datasets. Nat Commun *10*, 5415. 10.1038/s41467-019-13055-y.
5. Bolger, A.M., Lohse, M., and Usadel, B. (2014). Trimmomatic: a flexible trimmer for Illumina sequence data. Bioinformatics *30*, 2114-2120. 10.1093/bioinformatics/btu170.
6. Dobin, A., Davis, C.A., Schlesinger, F., Drenkow, J., Zaleski, C., Jha, S., Batut, P., Chaisson, M., and Gingeras, T.R. (2013). STAR: ultrafast universal RNA-seq aligner. Bioinformatics *29*, 15-21. 10.1093/bioinformatics/bts635.
7. Liao, Y., Smyth, G.K., and Shi, W. (2014). featureCounts: an efficient general purpose program for assigning sequence reads to genomic features. Bioinformatics *30*, 923-930. 10.1093/bioinformatics/btt656.
8. Team, R.C. (2022). R: A Language and Environment for Statistical, Computing
9. Love, M.I., Huber, W., and Anders, S. (2014). Moderated estimation of fold change and dispersion for RNA-seq data with DESeq2. Genome Biol *15*, 550. 10.1186/s13059-014-0550-8.
10. Zhou, Y., Zhou, B., Pache, L., Chang, M., Khodabakhshi, A.H., Tanaseichuk, O., Benner, C., and Chanda, S.K. (2019). Metascape provides a biologist-oriented resource for the analysis of systems-level datasets. Nat Commun *10*, 1523. 10.1038/s41467-019-09234-6.
